# A dual Purkinje cell rate and synchrony code sculpts reach kinematics

**DOI:** 10.1101/2023.07.12.548720

**Authors:** Abdulraheem Nashef, Michael S. Spindle, Dylan J. Calame, Abigail L. Person

## Abstract

Cerebellar Purkinje cells (PCs) encode movement kinematics in their population firing rates. Firing rate suppression is hypothesized to disinhibit neurons in the cerebellar nuclei, promoting adaptive movement adjustments. Debates persist, however, about whether a second disinhibitory mechanism, PC simple spike synchrony, is a relevant population code. We addressed this question by relating PC rate and synchrony patterns recorded with high density probes, to mouse reach kinematics. We discovered behavioral correlates of PC synchrony that align with a known causal relationship between activity in cerebellar output. Reach deceleration was positively correlated with both Purkinje firing rate decreases and synchrony, consistent with both mechanisms disinhibiting target neurons, which are known to adjust reach velocity. Direct tests of the contribution of each coding scheme to nuclear firing using dynamic clamp, combining physiological rate and synchrony patterns ex vivo, confirmed that physiological levels of PC simple spike synchrony are highly facilitatory for nuclear firing. These findings suggest that PC firing rate and synchrony collaborate to exert fine control of movement.

## Introduction

Cerebellar output is critical for online control of movement and cerebral cortical dynamics^1–6^. Purkinje cells (PCs) are the dominant input to neurons of the cerebellar nuclei and through their high spontaneous firing rates and inhibitory synapses place cerebellar output under constant inhibitory control. PC firing rate suppression is assumed to release nuclear cells from inhibition to influence behavior. However, a second disinhibitory mechanism may facilitate nuclear firing – synchronous PC activity – by reducing temporal integration of postsynaptic inhibition. Yet whether simple spike synchrony is a relevant mechanism for PC information transfer in vivo remains debated, with evidence for and against this mechanism playing an important role in cortical computations^7,8^.

While numerous studies have shown evidence for millisecond timescale PC simple spike synchrony, compelling PC rate codes have overshadowed synchrony as the dominant mechanism of information transmission^9–18^. Two recent studies in monkeys have renewed interest in this debate^7,8^. These groups built on an extensive literature on PC rate encoding of saccades and smooth pursuit kinematics, examining whether PC synchrony was also present during eye movements. During saccades in marmosets, synchrony between neighboring PCs in the oculomotor vermis increased markedly as the eye decelerated towards endpoint^8^, which, given previous studies could disinhibit the cerebellar nuclei and decelerate the eye^19,20^. In macaque flocculus, however, PC synchrony is minimal during smooth pursuit eye movements, and it was argued that synchrony may not be a behaviorally relevant code^7^. The source of these discrepancies is unresolved, but possibilities include regional, behavioral and analytical variations.

A particularly salient difference between saccades and smooth pursuit movements is that saccades require rapid acceleration and deceleration of the eye to achieve point-to-point movement, while smooth pursuit is dominated by steady-state eye velocities. Such differences differentially involve cerebellar computation^21,22^. If this difference contributes to the reported discrepancy, other point-to-point movements such as reaching would be predicted to engage synchrony. We therefore leveraged a previously collected dataset of simultaneously recorded PCs and putative PCs during mouse reaching movements to examine both the presence and relevance of physiological synchrony amongst populations of PCs in this behavior. Moreover, we took advantage of previous work showing that firing in the anterior interposed nucleus scales the decelerative phase of limb movements^23^, thus nuclear neuron activity can be inferred from kinematic variations.

We found that during reach synchrony increases markedly among PCs with behaviorally-associated firing rate suppression. This synchrony also related to movement kinematics: as the synchrony increased, the limb tended to decelerate faster and undershoot the target. Interestingly, synchrony and rate modulation tended to occur together, with both codes associated with kinematic variables. Finally, we established sufficiency of these PC population activity patterns on nuclear firing in ex vivo dynamic clamp experiments. Our results suggest that both rate and temporal modulations of the PCs complement each other to finely control the endpoint position of the limb, adding to the recent literature to suggest that the synchrony of PCs is an important mechanism by which the cerebellum controls fine movements.

## Results

### Purkinje cell synchrony increases during reaching behavior

We previously reported that mouse PC populations recorded *in vivo* show a net simple spike suppression that scales with reach velocity^10^, inverse to the code recorded in the cerebellar nuclei^23^. Similar patterns of PC activity have been observed in oculomotor vermis during saccadic eye movements, where it was also seen that PC simple spike activity synchronizes during movements. If these mechanisms are generalizable across regions of cerebellum mediating rapid, discrete movements, we might expect that PC synchrony would also be present in PCs mediating reach. To test this, we examined whether synchronous PC simple spikes are prevalent *in vivo* during mouse reaching movements. We recorded PCs and putative PCs (together referred to as pPCs; see Methods) using high density extracellular probes in forelimb associated lobules 4/5 and Simplex of cerebellar cortex^24^, during skilled reaching movements in head-fixed mice (same dataset used Calame et al^10^). We first analyzed whether synchrony of simple spikes occurred between simultaneously recorded pairs of identified and pPCs in reaching mice (Fig 1a) by calculating the joint peri-event time histograms-jPETHs^2,25,26^ between different populations at 1ms time bins. We divided pPC populations into groups of neurons that decreased their firing during reach (decreasers) and cells that increased their firing during reach (increasers; Fig 1b). After aligning to reach endpoint, jPETHs revealed prominent ms-timescale synchrony between pPCs that decreased their firing, with an uptick that preceded the endpoint and continued as a prominent signal afterward. pPC synchrony was modest or absent in pPCs that increased their firing rates as well as between those that increased or decreased rates during reach (Fig. 1c, f).

**Figure 1:**
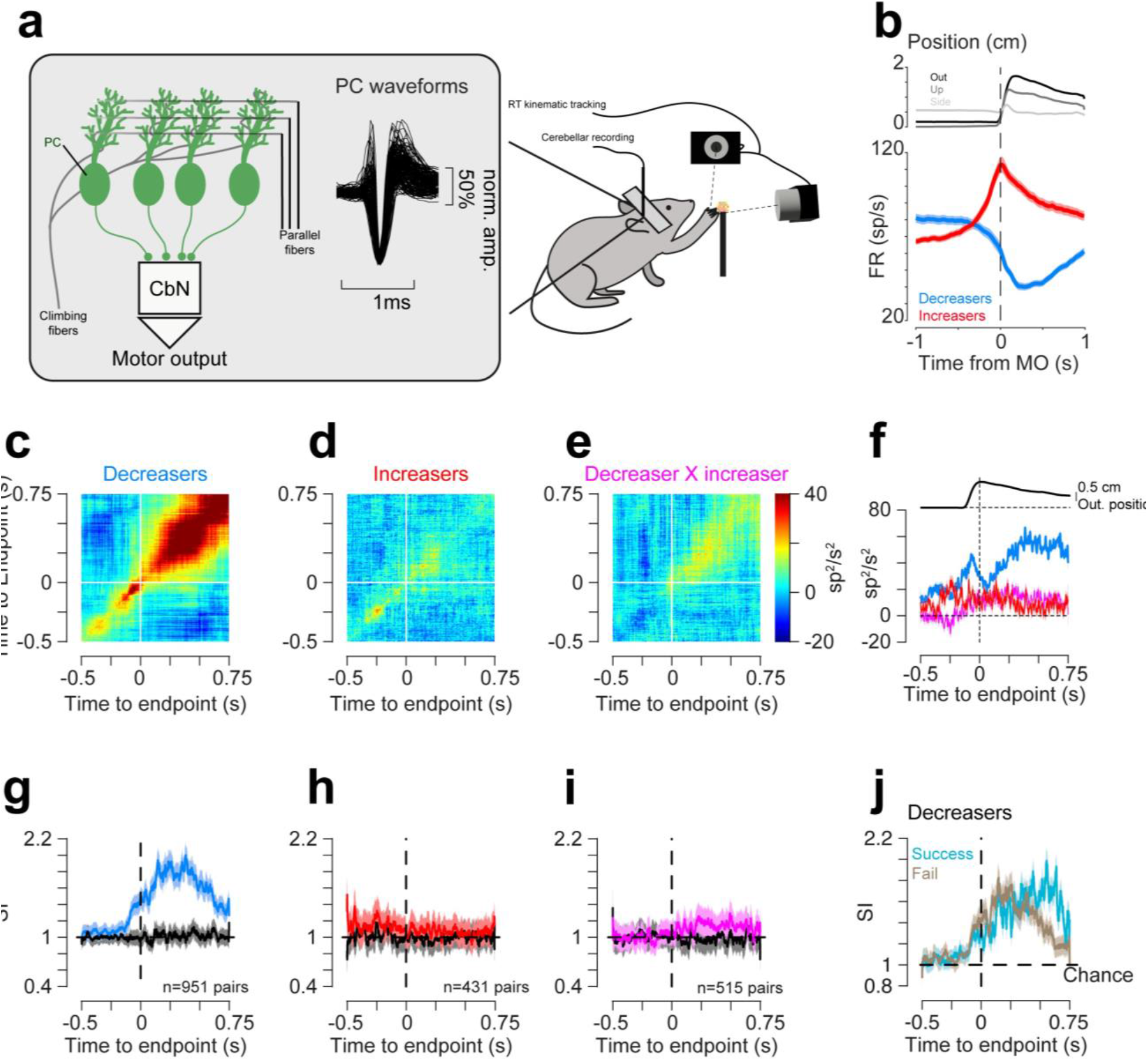
Dynamics of simple spike synchrony during reaching. (a) In head-fixed mice, neuropixels recordings were made from Lobules 4/5 and simplex while the mice performed reaching towards food pellets. Right panel shows the average normalized waveforms for units classified as Purkinje cells. (b) Top: average hand position in the outward, upward and lateral axes around time of motion onset (dashed line). Bottom: average (±s.e.m) firing rates of PCs/putative PCs (pPCs) around time of threshold crossing. Blue trace shows the average firing for the decreasers and red for the increasers. Shaded areas are the standard error of the mean (s.e.m.). (c) Joint peri-event time histogram (jPETH; see methods) calculated between the decreasers. Straight white lines indicate the time of endpoint; n=951 pairs. (d) Same as c, for increasers, n=431 pairs. (e) Same as c, for mixed pairs, n=515 pairs. (f) Matrix diagonals from jPETHs in c-e summarize the coordinated firing across the reach. Blue-Decreasers, red-increasers and magenta-mixed. The black line in top panel corresponds to the hand outward position. (g) Synchrony index (SI) between simultaneously recorded Purkinje cells during reaching around time of endpoint (dashed line). Black lines indicate the SI calculated when the trials were shuffled and blue line is the average SI for the real data, shaded areas around traces correspond to s.e.m.. (h) similar to g for increasers (red). (i) similar to g, but for pairs consisting of one decreaser and one increaser (magenta). (j) The SI between decreasers for either successful (blue) or failed (gray) trials around endpoint position.

To simplify comparisons with an intuitive metric of synchrony, we calculated the synchrony index (SI^8^), defined as the ratio of the observed probability of coincident simple spikes between two cells and the null probability given each cell’s firing rate (See Methods). Above-chance synchrony returns SI>1, while spike trains with independent spike times return a value of 1. In alignment with inferences from the jPETHs, we found a significant increase in SI in pPCs with suppressed firing rates around the time of endpoint position and afterward (n=951 pairs; Fig. 1g), in all animals tested (7/7) (Extended Data Fig 1a; 25% increase, p<0.05 compared to shuffled data). On the other hand, pPCs that increased firing rates during reach (n=431 pairs), and pairs of pPCs showing mixed responses (increases and decreases in firing rate, “mixed”; n=515 pairs) did not show a statistically significant increase in SI before or during the movement (Fig. 1h-i and Extended Data Fig 1b,c; Increasers: 5/6 animals; 22% increase; p=0.1. Mixed: 3/6 animals; 1.6% increase; p=0.52). Importantly, these patterns were also observed in confirmed PCs (Extended Data Fig. 1d-f). Among the decreaser population, the timing of synchrony increases suggested involvement in reaching and food handling behavior. Indeed, when we separated successful and failed trials we noted that elevated SI values following endpoint were driven by increased synchrony after successful reaches (Fig. 1j).

Although jPETHs and SI metrics aligned, a recent critique of the SI index as a measure of synchrony prompted a number of controls^7^. First, if we quantify the total number of synchronous events, without normalizing for ongoing firing rate, (i.e. remove the denominator from the SI equation), we still observe that the real simultaneously recorded trials have increased co-firing probability relative to shuffled controls, although the net number of coincident events drops, as is expected by the decrease in overall firing rate (Extended Data Fig 2). Second, if the SI in physiological signals is mainly driven by the underlying firing rate, we would expect the opposite behavior in SI with bursting activity causing drops in SI during the movement, which was not the case (Fig 1h). Third, we constructed artificial spike trains that mimicked firing rate suppression seen in the “decreaser” population. We then calculated the SI for the simulated data. As expected for independent spike trains, SI did not increase simply because firing rate drops (Extended Data Fig. 3). However, artificially adding synchrony to our simulated data in the time window between 0 and 0.5 s, increased the SI as the probability of synchronized spikes increased (Extended Data Fig. 3). Finally, if the SI is merely an artifact of firing rate in trains that have covariance, we would expect that the shuffling procedure we used, which preserves firing rate fluctuations, would have resulted in SIs that increase, but this was not seen. Therefore, we conclude that during reach, pPCs with suppressed firing rates selectively synchronize simple spikes. Given the modest and non-significant SI modulations for increasers (and mixed) populations, we focus our analyses below on neurons that decreased their firing during reaching.

**Figure 2:**
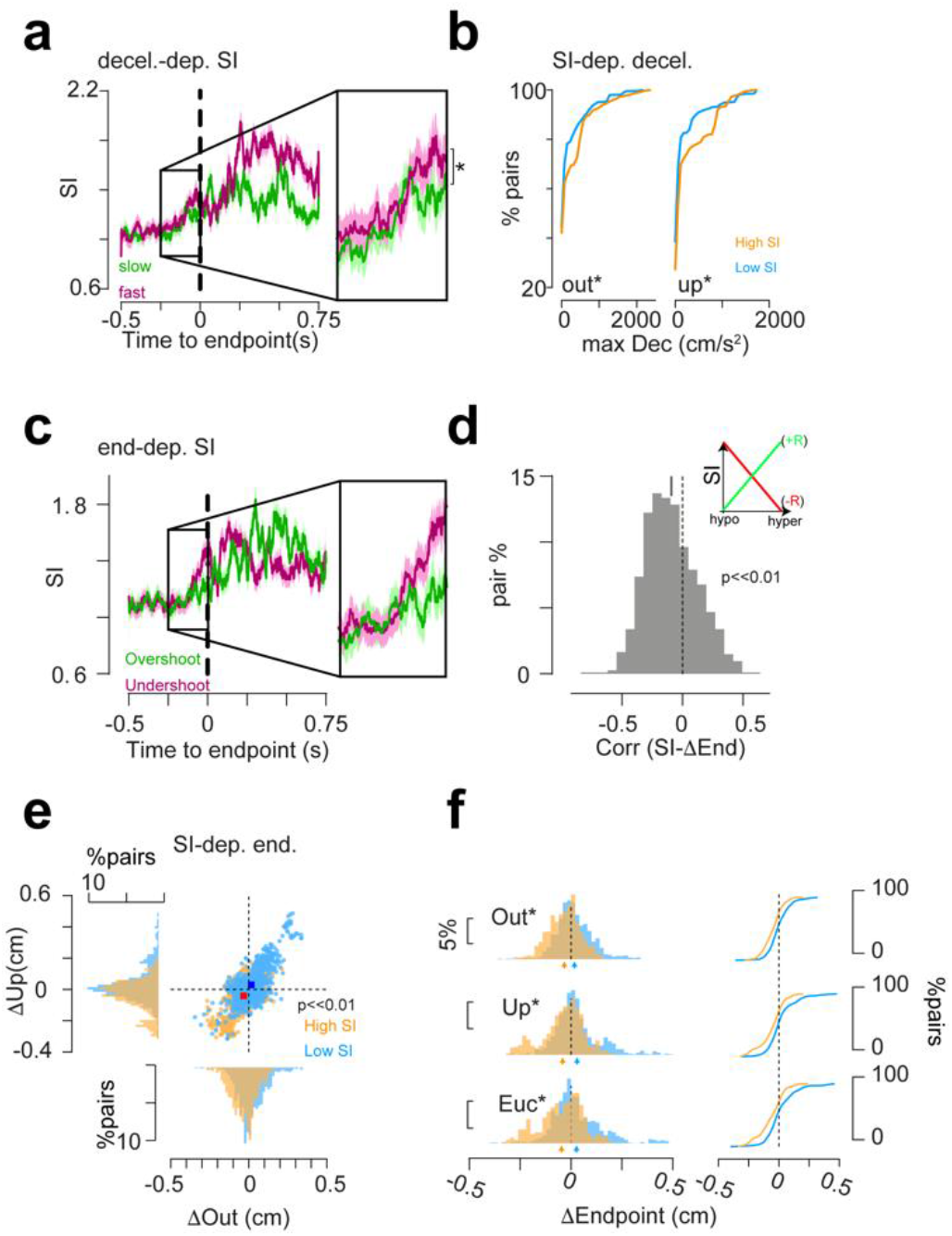
Relationship between SI and behavioral kinematics. (a) For each session, the SI was calculated between different pairs for trials with fastest (green) or slowest (purple) decelerations.Insets show magnification of the difference between the two conditions in the time leading to endpoint (Average (±s.e.m.) SI in times −0.3 to 0s around the endpoint; Slow: 1.07± 0.02; Fast: 1.13±0.02; Wilcoxon’s signed rank; p=0.02; r=0.08). (b) The maximal deceleration observed for trials with the highest SI (orange; calculated for the 0.3 s preceding endpoint time) or trials with the lowest SI (blue) in the outward (Out.) or vertical (Up.) position (n=951 pairs. Out.: Low SI: 227.2±11.9 cm/s^2^; High SI: 316.5.4±14.4 cm/s^2^; Kolmogrov-Smirnov test: p=5.7×10^-9^; Up.: Low SI: 201.5±11.5 cm/s2; High SI: 297.8±12.7 cm/s^2^;p=6.2×10^-14^). (c) For each session the SI was calculated between different pairs for trials with hypermetric (overshoot; green) or hypometric (undershoot; purple) reach endpoints. Points on top correspond to time points where the two traces have significant difference (p<0.05). Inset shows magnification of the difference between the two conditions in the time leading to endpoint. (d) Correlation between the sum of the SI per trial and the endpoint changes (mean±s.e.m.; −0.09±0.007; p=4.2×10^-35^; r=0.4). (e) Endpoint locations for trials with the highest SI (orange), and trials with the lowest SI (blue), per pair of cells (n=951 pairs; Low SI: Δout: 0.016±0.003 cm; Δup: 0.03±0.004 cm; High SI: Δout: −0.033±0.003 cm; Δup: −0.04±0.003 cm; Wilcoxon’s signed rank; p=1.2×10^-48^; r=0.48). Also shown is the distribution of the outward (bottom) or upward (left) endpoint position per condition. (f) The change in the outward (Out.), vertical (Up.) or Euclidean (Euc) endpoint position for low SI (blue) and high SI (orange) trials (Kolmogrov-Smirnov test; Δout: p=1.4×10^-21^; Δup: p=5.4×10^-21^; Euc.: Low SI: 0.027±0.004, High SI: −0.047±0.003, p=8.2×10^-24^).

**Figure 3:**
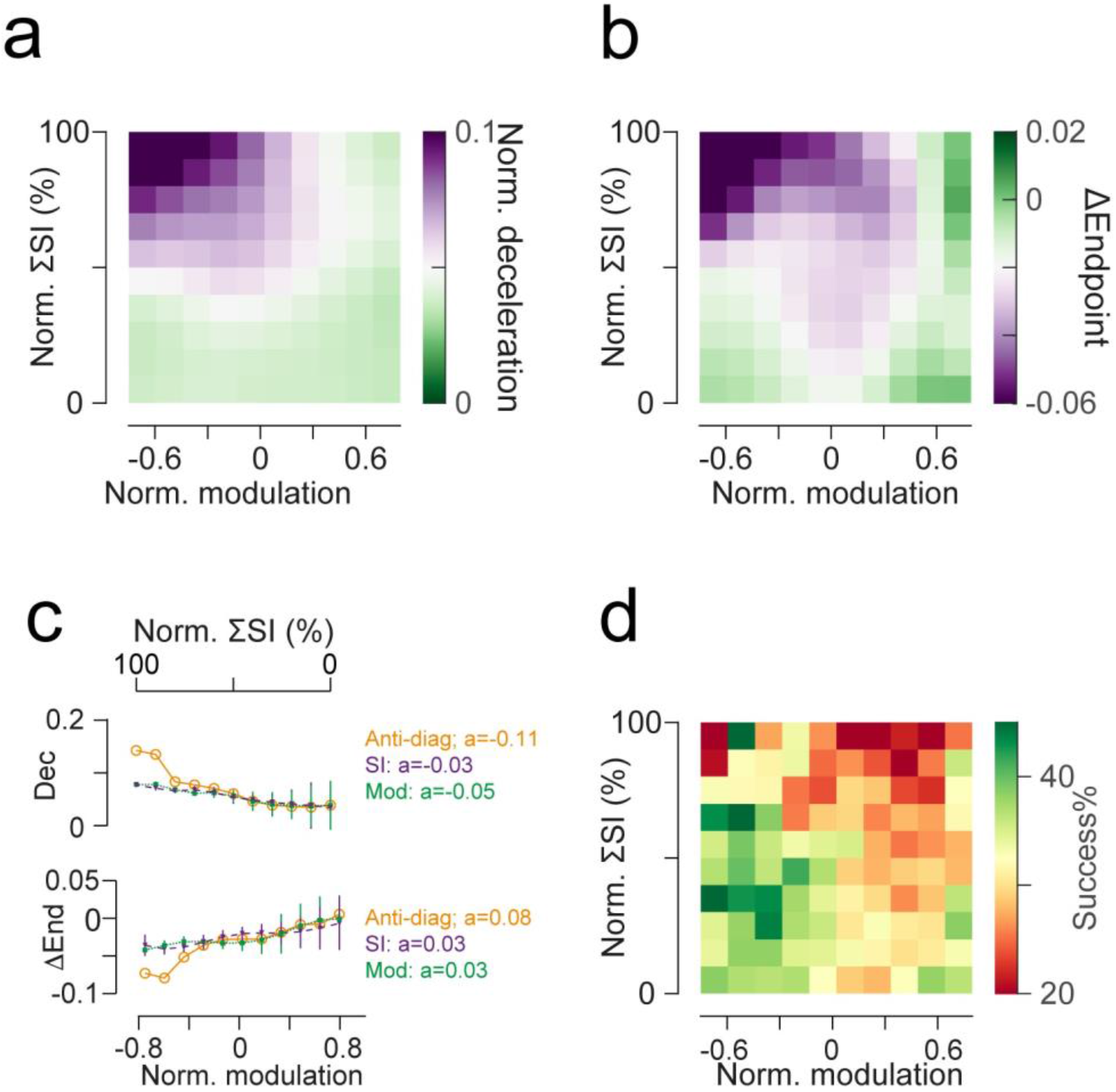
Synchrony and rate modulation jointly influence movement kinematics. (a) Heatmap showing the average deceleration as a function of SI (y-axis) and modulation changes (x-axis). The SI and Modulation were calculated for time window of −0.3 to 0.3 s around the endpoint position to capture the whole movement and normalized on a session basis. The deceleration was also normalized per session to avoid behavioral variability between sessions and mice. (b) Same as (a), for the relationship between the SI, the modulation and the reach endpoint, difference in endpoint position was calculated as the change in trial-by-trial endpoint position from the average endpoint for a session. (c) Change of deceleration (dec, top) and endpoint (end; bottom) as function of changing Modulation (green), changing SI (purple; Note that the SI axis is reversed) or both (calculated from the antidiagonal of the matrices in a-b (orange. The slope of each scatter is shown. (d) Same as a, for the relationship between the SI, rate modulation, and reach success rate.

### Kinematic correlates of PC synchrony

The data above are consistent with previous findings showing increased synchrony during saccadic eye movements in monkeys^8^. If synchrony is a relevant component of Purkinje population coding, however, we would predict that there should be behavioral consequences of synchrony. Previous studies strongly constrain predictions on how pPC simple spike synchrony in this region would influence reaching movements: First, in vitro studies show that increased synchrony promotes nuclear firing^15^. Second, increased activity in IntA during reaching decelerates the limb on outreach^23^. Thus, we would predict that increased synchrony during the reach would promote more nuclear firing which in turn would exert a stronger inward pull on the limb, leading to faster deceleration and hypometric endpoints. To test this behavioral prediction, we sorted reaches by deceleration and compared SI for grouped trials (fast deceleration (top quartile) vs. slow deceleration (bottom quartile)). We found that trials with faster deceleration tended to have higher SI before endpoint (Fig 2a). Additionally, by calculating the trial-by-trial relative SI (see methods), we found that deceleration was faster on reaches with higher SI, compared to low-SI trials (Fig. 2b; n=951 pairs. Out.: Low SI: 227.2±11.9 cm/s^2^; High SI: 316.5.4±14.4 cm/s^2^; Kolmogrov-Smirnov test: p=5.7×10^-9^; D= 0.14; Up.: Low SI: 201.5±11.5 cm/s^2^; High SI: 297.8±12.7 cm/s^2^;p=6.2×10^-14^; D= 0.18).

In addition to deceleration, increased activity from anterior interposed can promote hypometric reaches. Therefore, we next examined the relationship between SI and hand endpoint position (Fig. 2c-f; defined as maximal outward extent). We found that hypometric reaches (undershooting; average Δout=-0.28 cm, ΔUp=-0.19 cm) tended to have higher SI before endpoint, compared to hypermetric reaches (overshooting; Δout=0.12 cm, ΔUp=0.07 cm; Fig 2c). In addition, the trial-by-trial relative SI was negatively correlated with the endpoint position (mean±s.e.m.: r = −0.09±0.01; Wilcoxon’s signed rank p=4.2×10^-35^ against 0; Fig 2d). In a converse analysis, we sorted trials by SI, and examined endpoint position. Trials with the highest SI had the nearest endpoints (endpoint calculated relative to the session averaged endpoint position to avoid behavioral differences between days and mice). This showed that for trials with the highest SI, the movement tended to be hypometric, and vice versa (ΔOut- Low SI: 0.016±0.003 cm; High SI: −0.033±0.003 cm, p=3×10*^-25^*; ΔUp- low SI: 0.03±0.004 cm; high SI: − 0.04±0.003 cm, p=3.3×10*^-25^*; ΔOut-Up: p=1.3×10*^-48^*; Euc.: Low SI: 0.027±0.004, High SI: − 0.047±0.003, p=1.3×10*^-29^*; Wilcoxon’s Signed rank; Fig. 2e-f). Importantly, cells with identified complex spikes also had a similar relationship (Extended Data Fig 4). A similar relationship between co-firing and endpoint position was found (Extended Data Fig. 5). These data also suggest that the prolonged elevation of SI seen in successful reaches (Fig. 1j) may support limb flexion associated with food consumption.

**Figure 4:**
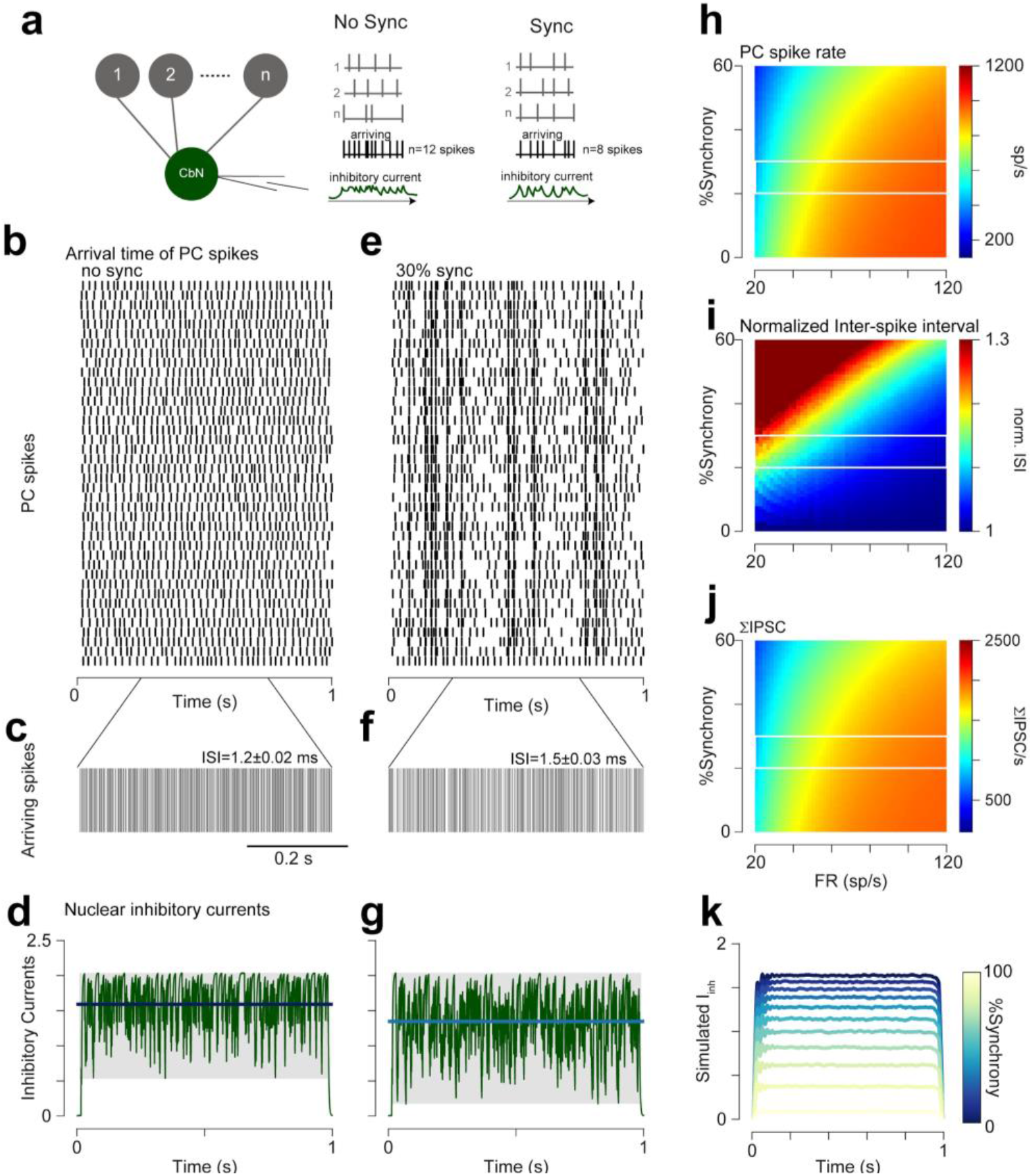
Nuclear response to PC firing simulation. (a) schematics of the model. Nuclear cell (Nu) receive input from n PCs (n=40). When there is not synchrony, the number of spikes that arrive at the Nu is the same as the total number of spikes for all the converging PCs (“arriving”), however, when synchronized, spikes superimpose, such that >1 spikes can arrive at the same time at the Nu, eliciting short and one IPSC, such that the total number of arriving spikes is lower than that of the total spike number that was fired by the PCs. (b) example of non-synchronized PC firing, for 40 cells. (c) the train of spikes that arrive at the CbN, each line correspond to ≥1 spikes at the specific time bin. The average inter-spike interval (ISI) value ± s.e.m. is shown above the trace (d) the simulated Nu inhibitory currents elicited by the PC spikes. Each spike elicited an IPSC with normalized amplitude 1 and τ =2.5ms (40 sp/s firing rate). Bold line shows the average current and gray shaded area the range of currents observed. (e) Same as b, for 30% synchronized spiking. (f) same as c, for synchronized spiking. (g) same as d, for synchronized spiking (30% synchrony). (h) The number of spikes that arrive at the Nu, as a function of the synchrony % for different baseline firing of the PCs (n=40 converging cells). The data represent 1,000 iterations of the model. White lines show the range of synchrony levels that are observed in vivo. (i) The average inter-spike interval (ISI) observed by the Nu cell given the different FR and synchrony level. (j) The integral of the inhibitory currents (see d) as a function of increasing synchrony and different firing frequencies. (k) the average change in inhibitory current as a function of the synchrony increase for 40 converging PCs firing at 40 sp/s. Same as d and g, for 1,000 iterations.

**Figure 5:**
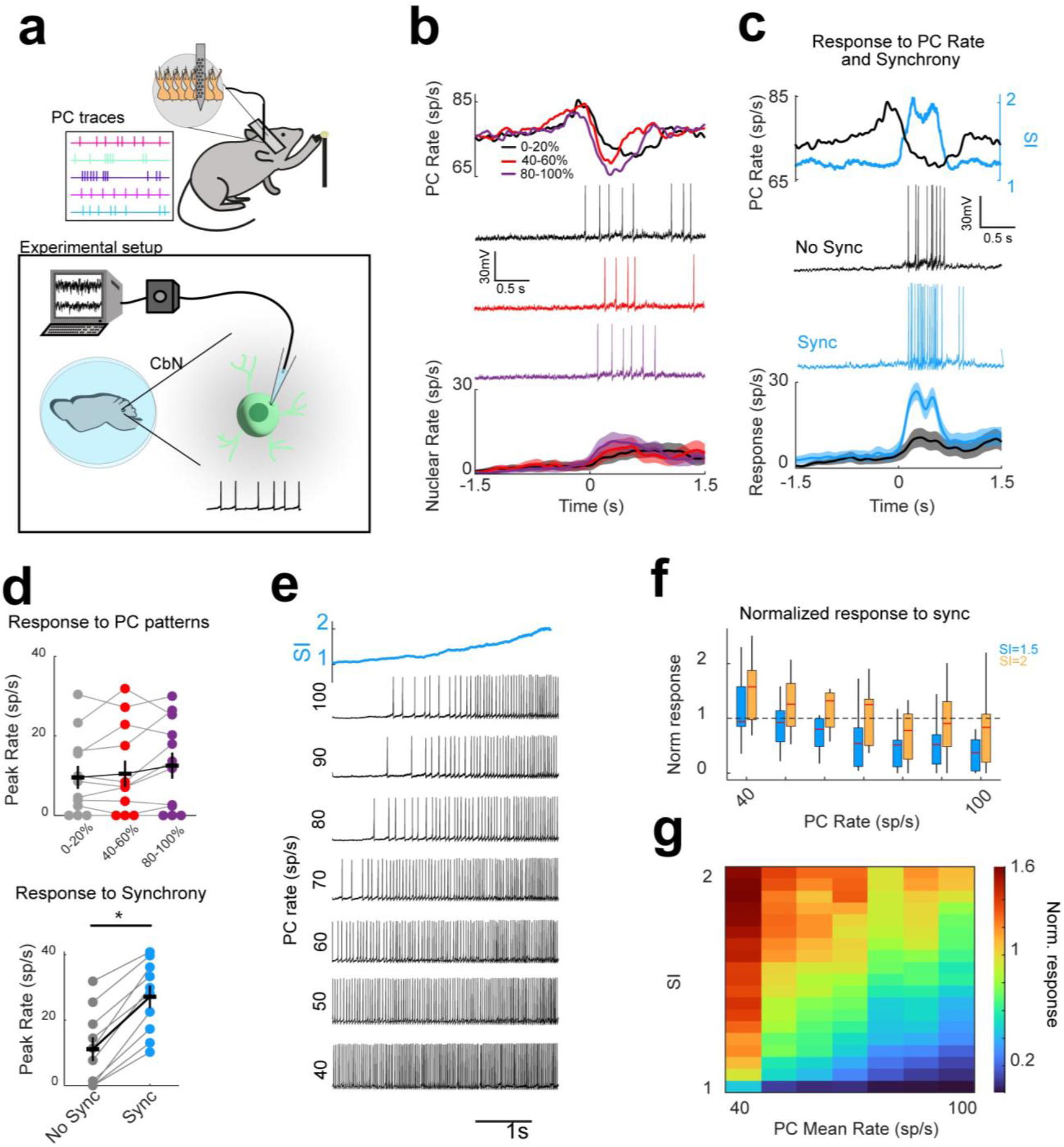
Synchrony and rate are required for eliciting nuclear firing in vitro. (a) Schematics showing the dynamic clamp experiment setup. (b) Nuclear response to simulated PC trains. From top to bottom: PC pool peri-event time histogram (PETHs) used to construct IPSCs trains (from Calame et al.^10^), example traces of nuclear cell response to each pool, and mean nuclear response (mean±s.e.m.; n=13 cells). (c) Nuclear cell responses to pooled PCs (black) with synchrony (blue) as derived from our in vivo measurements. Middle panel shows representative voltage traces for responses to non-synchronized (black) and synchronized (blue) PC pools. Bottom panel shows the mean nuclear response (± s.e.m.) to the PC activity with (blue) or without (black) synchrony. (d). Summary data for peak nuclear responses. Mean nuclear peak rate when the 2 PC trains had no applied synchrony (no sync; from b; 9.57±2.87, 10.5±3.32, and 12.6±3.28 sp/s, respectively, 1 way ANOVA F=0.23, p=0.79 compared to the nuclear response with applied synchrony (no sync:11.2±3.6 sp/s. sync: 27.1±3.5 sp/s; paired t-test: p=3×10^-6^; n=10 cells). (e) Representative traces of nuclear responses to PC pools held at a fixed mean rate in 10 sp/s intervals, while synchrony (blue trace) was increased throughout the trace. (f) normalized nuclear response to PC pools with a static rate between 40 and 100 sp/s, and SI of either 1.5 or 2 (n=10 cells; Two-ways ANOVA; F_6_(PC firing)=5.15, p=1×10^-4^; F_1_(SI)=15.62, p=1.4×10^-4^). (g) Heatmap showing the change in the nuclear response as a function of PC rate (x-axis) and SI (y-axis).

These data suggest that synchrony has behavioral consequences, consistent with a disinhibitory effect on the cerebellar nuclei.

### Synergistic contributions of PC rates and synchrony to kinematics

The observations above show a clear relationship between synchrony and reach kinematics. However, a vast literature shows strong relationships of pPC firing rates with kinematic variables^11,13,14,16,27–29^, including within the present dataset^10^, raising the question of whether both rate and synchrony may signal in tandem. To test whether pPCs may control reach endpoints using both coding mechanisms, we related kinematics to both the cumulative normalized SI over the reach epoch and the firing modulation relative to baseline. Visualizing relative reach kinematic variables as a function of both rate modulation (horizontal axis) and SI (vertical axis), (Fig. 3 and Extended Data Fig. 6), revealed a striking pattern: both deceleration and endpoint undershoots were most pronounced when both pPC rate suppression and SI were highest, suggestive of a synergistic disinhibitory code in PCs (Fig. 3a-b and Extended Data Fig.6). We quantified this synergy by comparing kinematic changes along the matrix antidiagonal (which captures both changing SI and pPC rates) to the averages of either the matrix rows (firing rate, agnostic to SI) or columns (SI, agnostic to firing rate), to examine the role of either one in the control of the reach (Fig. 3c). This analysis showed that together, the two codes are more strongly related to reach kinematics than either one individually, as indicated by the flatter relationship of each code and the movement (Linear regression slopes: Deceleration-Antidiagonal: −0.11; modulation alone: −0.05; SI: 0.03/ Endpoint-Antidiagonal: 0.08 modulation: 0.03; SI: −0.03; p<0.001 for all). This steeper relationship suggests that the kinematic changes are stronger when both variables change. To further interrogate this relationship, we next measured behavioral variability as a function of the two coding mechanisms, either individually or together. We fitted the bin-by-bin values (Fig. 3a-b) of SI and rate modulation to the kinematic data using multiple regression models (see methods). Regressing kinematics to SI and rate modulation together accounted for more variance than either code individually (Multiple linear regression model; Deceleration: Dual: R^2^=0.62, F_118_=99.5; SI-alone: R^2^=0.33, F_119_=58.7; Rate-alone: R^2^=0.29, F_119_=50.4/ Endpoint: Dual: R^2^=0.55, F_118_=72.9; SI-alone: R^2^=0.21, F_119_=33.3; Rate-alone: R^2^=0.33, F_119_=59.6; p<<0.001 for all). Together, these results indicate the relevance of both rate and synchrony modulation in the control of reach kinematics.

The extremes of the SI:firing rate relationships (high SI, deep pPC inhibition or low SI and increased pPC rate) were associated with hypo- or hypermetric reaches. We therefore wondered if there were particular patterns of pPC rate and synchrony associated with successful reaches, when the mouse acquired the food pellet. Analyzing the likelihood of successful trials given the rate modulation and the SI, we observed that successes were most probable with deep rate suppression but moderate synchrony (Fig. 3d, dark green regions), while maximal and minimal synchrony were more associated with failures (Fig. 3d, red regions). Thus, these data suggest that cerebellar cortical mechanisms may exist to finely regulate these parameters in tandem, rather than maximize either. To summarize, synchrony does not necessarily lead to more successful trials, yet rate and synchrony together both influence movement kinematics.

### Mechanisms of PC synchrony on regulating nuclear output

Synchronous firing in the cerebral cortex facilitates information transfer through strong temporal summation of excitatory synaptic potentials, driving postsynaptic firing with greater efficacy^30,31^. By contrast, inhibitory PC synchrony facilitates nuclear neuron firing by effectively reducing the summated firing rate of convergent PCs. While this mechanistic intuition has been articulated previously^3215^, the quantitative relationship of PC population synchrony and the “effective input rate” (i.e. total IPSCs/s minus co-occurring IPSCs) to the nuclei, incorporating physiological convergence ratios, rates, and synchrony has not been explicitly defined. Thus, we built a simple model that simulated the firing rate of 40 PCs converging onto a nuclear cell (Fig. 4). Firing rates and synchrony were varied systematically. This model shows that as synchrony increases, the effective input rate to the nuclear cell decreases. Moreover, overlapping synchronous spikes result in longer inter-spike intervals within the convergent population (Fig. 4c,f,i), and less summated inhibitory current in the nuclear cell (Fig. 4d,g,j,k). Overall, physiological synchrony levels seen in vivo, which correspond to peaks ∼25%, are equivalent to a reduction in PC firing rates on the order of 5 spikes/s per cell, within the range of those seen with behaviorally relevant rate codes^10,13,16,33,34^.

### Synergistic contributions of PC rates and synchrony to nuclear disinhibition

The results above show consistent correlations between PC simple spike synchrony in the ‘decreaser’ population and behavioral metrics associated with nuclear firing. To determine the sufficiency of rate and temporal patterning of PC activity on nuclear firing using the full population of increasers and decreasers together, we turned to using dynamic clamp in acute adult mouse brain slices (Fig. 5). We assayed nuclear neuron responses to pools of 40 PCs, matching firing rate patterns recorded in vivo during reaching movements that varied in peak velocity and depth of firing rate suppression (0-20%; 40-60% and 80-100% of peak reach velocity^10^). Inhibitory postsynaptic conductances were modeled based on physiological constraints previously established^15^(See Methods). With no above-chance population synchrony, these naturalistic PC trains resulted in an almost complete suppression of nuclear neuron firing, with spontaneous firing rates inhibited to 2.34 ± 1.0 sp/s. As expected, PC suppression disinhibited nuclear neurons, promoting firing up to a mean of 11.5 ± 2.29 sp/s at their peak during the maximal disinhibitory phase (Fig 5b). Interestingly, however, nuclear firing rates were relatively insensitive to variations in PC input rates (Fig 5b,d; from b; 9.57±2.87, 10.5±3.32, and 12.6±3.28 sp/s, respectively, 1 way ANOVA F=0.23, p=0.79), and was minimal compared to nuclear firing rates recorded in vivo.

Next, we imposed synchrony within the naturalistic PC trains tested (Fig. 5c,d). Firing rates were dramatically increased with the combination of PC rate suppression when the synchrony index ranged from 1.5-2 – levels within the physiological range (Fig. 5c-d; 144.1% increase; no sync:11.2±3.6 spike/s. sync: 27.1±3.5 spike/s; paired t-test: p=3×10^-6^; n=10 cells). These data suggested that both rate and synchrony act synergistically to disinhibit nuclear firing. Finally, we explored the ‘synergy’ between PC rates and synchrony in driving nuclear firing, varying both rate and synchrony across a broad range of values. Both PC firing rate and PC synchrony influenced nuclear firing, with the strongest disinhibition occurring with low PC rates and high PC synchrony, and the strongest inhibition with the converse pattern, high PC rates and low PC synchrony (Fig. 5e-g; n=10 cells; Two-ways ANOVA; F_6_(PC firing)=5.15, p=1×10^-4^; F_1_(SI)=15.62, p=1.4×10^-4^). Together, these data suggest that synchrony can influence the gain of PC:nuclear firing rates.

## Discussion

In this study, we found evidence that Purkinje cells in mouse intermediate cerebellar cortical lobules use two mechanisms simultaneously to adjust downstream activity to influence reaches. Subpopulations of PCs in a limb-related region show higher-than-chance probability of synchronous spikes selectively during reaching movements as well as reduction in overall firing rates. Moreover, ex vivo dynamic clamp experiments showed little evidence of effective disinhibition from physiological PC rates alone, but strong responses when physiological patterns were coupled with synchrony. In keeping with this observation, kinematic variables of reach were correlated with both rate modulation and synchronous firing. Both synchrony and rate suppression correlated with limb deceleration and hypometric reach endpoints, consistent with the previously established role of the cerebellar interposed nucleus regulating reach. These results suggest that populations of PCs, at least in the region of intermediate cerebellum, use both firing rate and synchrony to regulate nuclear firing and movement kinematics, ensuring rapid and precise control.

Millisecond-level synchrony between PCs has been widely observed in vivo across multiple species and conditions, yet whether synchrony plays a behaviorally relevant role has remained hotly debated^7–9,12,17,18,35–39^. In behaving animals, PC rate codes strongly correlate with both downstream (postsynaptic) rates and behavior, casting doubt on the importance of a secondary synchrony code^7^. In other behaviors that involve distinct cerebellar regions, however, synchrony appears concomitant with well-timed behavioral events such as whisker deflections and saccades^8,37,38^. Several differences may underlie these discrepancies. Regional specializations across cerebellar regions exist, with distinct distributions of unipolar brush cells, zebrin striping, density of Golgi cells, and cerebello-olivary interactions^40–43^. These circuit motif variations may lead to distinct computations supporting behaviors with diverse control system requirements. Smooth pursuit eye movements, which require the floccular complex, are non-ballistic, thus inertial non-linearities are minimal. By contrast, saccadic eye movements, which are supported by the oculomotor vermis, require accelerative and decelerative signals to overcome inertial forces. It is possible that these or other unique control requirements require distinct neural mechanisms across cerebellar regions. In line with this reasoning, the present data corroborate Heck et al.^12^, finding evidence for behaviorally modulated synchrony during reaching movements, which also require marked control of accelerative and decelerative forces.

A limitation to the previous studies is a dearth of information about the mapping of cerebellar output neurons – the targets of PCs – to behavioral variables. The present work leveraged previous discoveries that the anterior interposed exerts an inward pull on the limb during reach, with firing rates scaling the rate of outward deceleration^23^. Additionally, PC neurons that project to this region show an inverse rate code in the population of PCs, with net PC suppression scaling with reach velocity (and therefore deceleration^10^). This knowledge constrained predictions about the effect of PC firing rates and synchrony on the behavior: any code associated with faster deceleration and hypometric reach endpoints is likely to map onto increased firing in the cerebellar nuclei. Because the synchrony we observe was coupled to PC rate suppression and enhanced reach deceleration leading to hypometric reach endpoints, it is strongly suggestive of an amplified disinhibitory effect on the cerebellar nuclei. In particular, this finding is consistent with previous observations that activity in the interposed nucleus regulates reach precision by braking reaching movement^23,44,45^.

In cortical motor areas, neural synchrony during behavior is a well-studied mechanism by which different brain areas influence movement^2,46,47^. Temporal coherence within 1 ms in sensory and motor areas of the neocortex is seen selectively during movements or to distinct sensory stimuli^48–50^. While synchrony between excitatory projection neurons can enhance temporal summation to threshold in downstream neurons^51^, synchrony between inhibitory projection neurons, by analogy, might be expected to enhance inhibition of postsynaptic cells. The opposite is seen, however, between PCs and cerebellar nuclei, where synchronous PC firing has a strong ‘disinhibitory’ effect on cerebellar nuclear neurons in brain slices^15,32,52–54^ and in vivo when PC synchronization is artificially induced^55^. The underlying cause of this disinhibitory effect is reduced inhibitory current accumulation over time and the associated enhancement of inhibitory current variance. Thus, PC synchronization is equivalent to reduced firing rates at the population level and may thus modulate the gain of PC-to-nuclear rate relationships when combined.

Conversely, asynchronous IPSCs will more effectively inhibit nuclear firing by accumulating ‘tonic’ current, thereby reducing the gain of PC-to-nuclear rate relationships. We found that physiological-levels of synchrony reduce the ‘effective’ rate of Purkinje populations impinging on nuclear neurons, equivalent to the PCs dropping their firing rates by 5-10 spikes/second, similar to rate changes seen with rapid PC-dependent learning^34^. Importantly, our model shows that this is PC rate-dependent, with enhanced disinhibition from synchrony when the PCs are least active.

The presence of synchrony in mouse and monkey Purkinje populations – at least in some cerebellar regions – raises the important question of the circuit mechanisms that promote it. Many proposed mechanisms have received empirical support, including shared parallel fiber afferents^12^, shared climbing fibers^18^, shared MLI inputs^56^, as well as ephaptic and gap junctional coupling between cerebellar cortical cell types^39,57,58^. The latter phenomena have been shown to synchronize MLIs which, through feedforward inhibition, regulate spike-timing and synchrony in PCs by shortening integration time windows for coincident excitation^57–63^. The relationship between MLIs and synchrony is particularly intriguing given the enhancement of PC co-firing during firing rate suppression that we and others have observed^8^. It will be interesting to investigate whether such circuit elements may be differentially organized in regions with or without regulated PC synchrony, or if mechanisms exist that explicitly degrade PC synchrony.

Overall, our study presents new evidence indicating that PC synchrony works synergistically with firing rate to regulate downstream neuron activity and control movement. It will be interesting for future studies to investigate similar coding principles in long-range inhibitory neurons in other brain regions^64^, such as striatal and pallidal regions of the basal ganglia, noting that both healthy and pathological states may be influenced by synchronization^65–67^. Moreover, given the association of the highest synchrony with failed reaches and kinematic disruption, associating pathological synchrony with cerebellar disorders could be a fruitful avenue of investigation^68–71^. With the advent of dense recording methods, determining the generalizability of such processes is within reach.

## Extended Data Figures

**Extended Data Figure 1:**
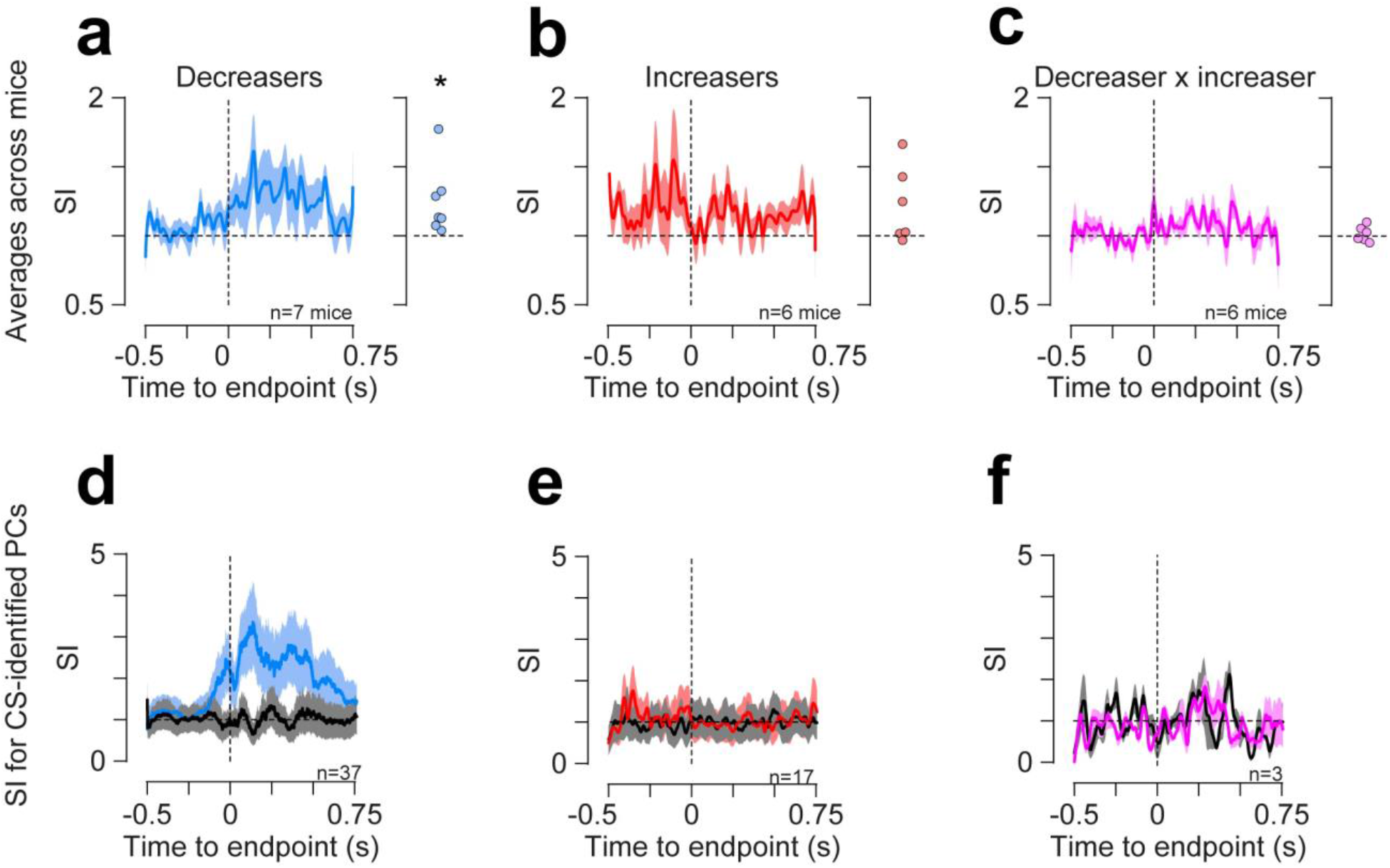
Synchrony index averaged across mice for all pPCs (a-c) and for confirmed PCs (d-f). (a) Left, average SI for decreasers calculated across 7 mice. Right, average SI in the time window between −0.3 and 0.3 s of the reach for each mouse (dots) (paired t-test; p<0.05 compared to shuffled data). (b) Same as a) for increasers (p=0.1). (c) Same as a) for mixed pairs (p=0.52). (d-f) SI of complex spike-confirmed PCs that decrease their firing (d;n=37 pairs) increase their firing (e;n=17 pairs) or decrease vs. increase pairs (f, n=3 pairs), relative to shuffled data (black).

**Extended Data Figure 2:**
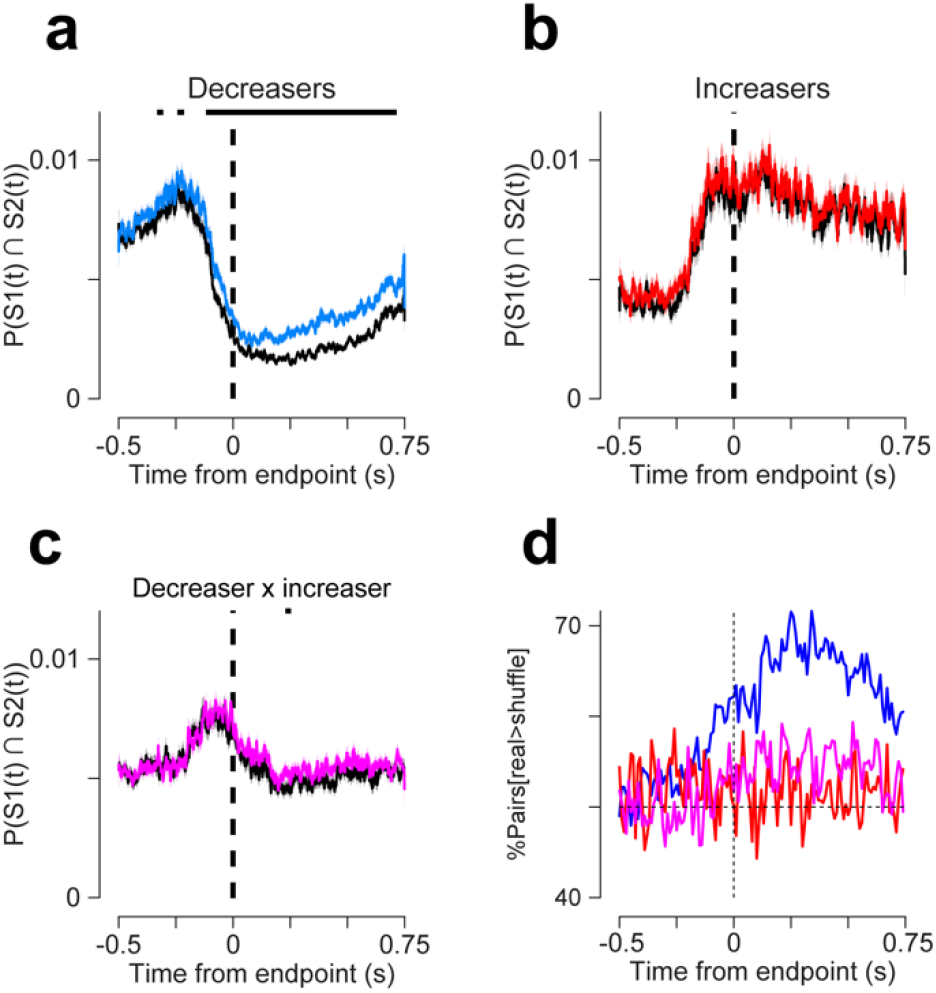
Probability of coincident spikes between pPCs. (a) Co-firing of pPCs across time relative to reach endpoint. Co-firing (colored traces) was calculated as the probability of sampling two spikes simultaneously from two cells in a 1-ms time bin (not corrected for average firing rate). Black curves report co-firing in shuffled data. Asterisks above show timepoints which were statistically significantly different from shuffle (paired t-test; p<4×10^-5^; Bonferroni corrected for multiple testing). (b) Same as a, but for increasers. (c) Same as a, but for pairs of increasers vs decreasers. (d) The percentage of pairs that exhibit higher co-firing relative to shuffled data, as a function of time from reach endpoint.

**Extended Data Figure 3:**
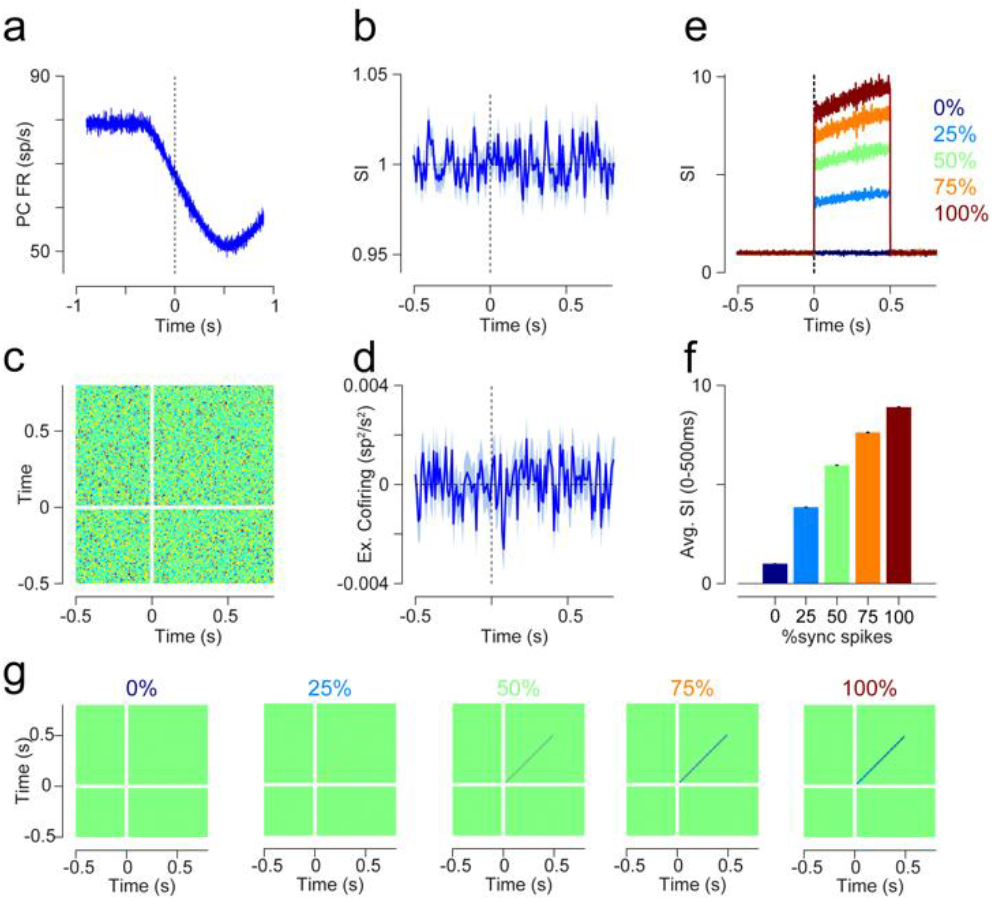
Data simulations to test sensitivity of synchrony index to firing rate. (a) Average simulated firing rate of simulated PCs that decrease rates. The simulation (see methods) was made for 500 pairs of decreasers, with 60 trials. (b) Average (±s.e.m.) synchrony index of the simulated data. (c) jPETH of the simulated data. (d) Average diagonal of the jPETH matrix in c. (e) The synchrony index after artificially synchronizing the firing of simulated pairs in the time window between 0 and 0.5 s. (f) The average SI (between 0 and 0.5s) from panel e (from 0% to 100% spike synchrony: 1±0.002; 3.9±0.01; 6±0.02; 7.6±0.02; 8.9±0.03). (g) From left to right, the jPETH for the different synchronization levels (0, 25, 50, 75 and 100%) in the time window between 0 and 0.5 s.

**Extended Data Figure 4:**
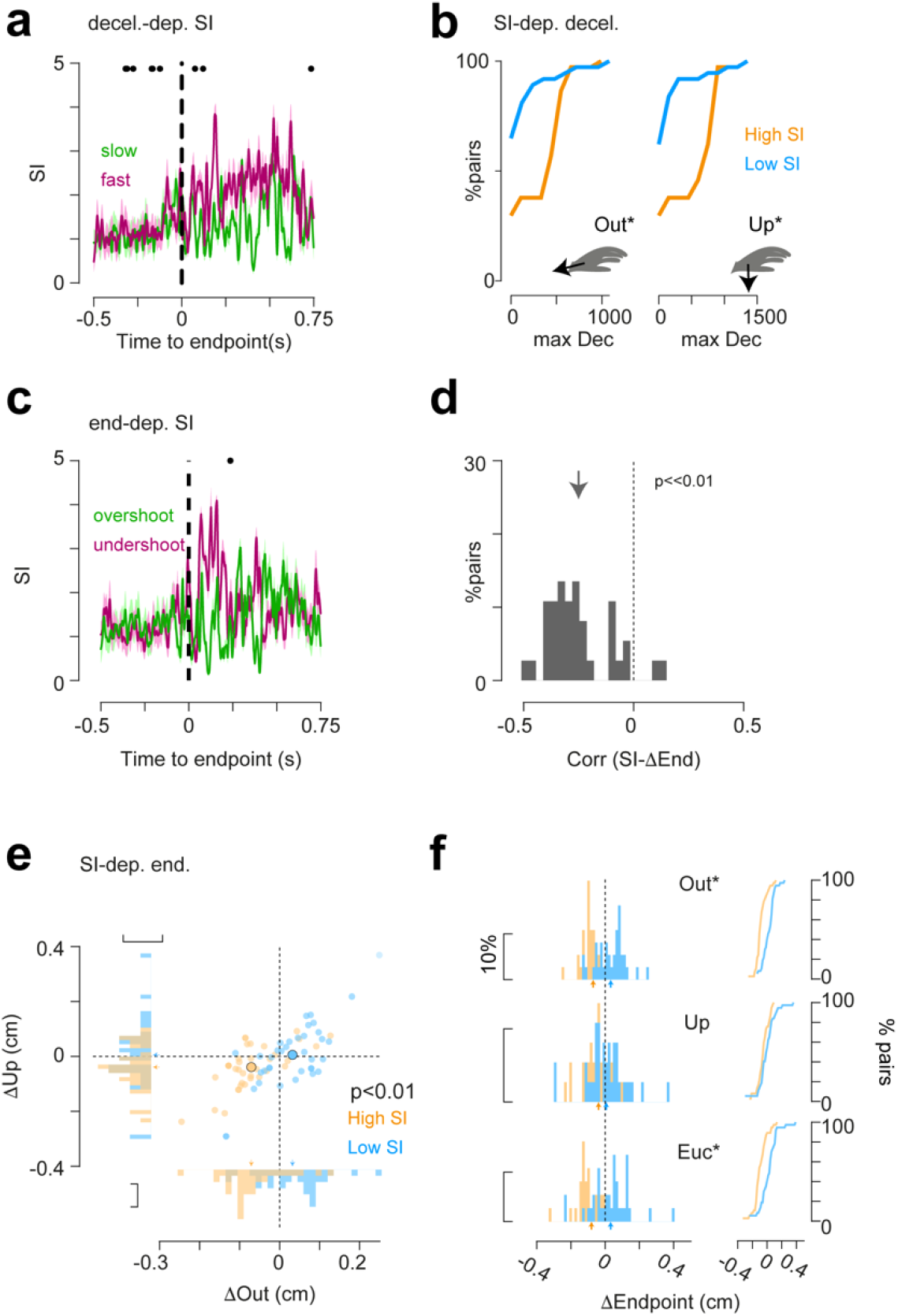
Relationship between synchrony and movement kinematics in confirmed PCs. (a) For each session the SI was calculated between different pairs for trials with fastest (green) or slowest (purple) decelerations. Points on top correspond to time points where the two traces have significant difference (p<4×10-5 Bonferroni corrected for multiple testing). Inset show magnification of the difference between the two conditions in the time leading to endpoint. (b) The maximal deceleration observed for trials with the highest SI (orange) or trials with the lowest SI (blue) in the outward (Out.) or vertical (Up.) position (n=37 pairs. Out.: Low SI: 177.4±38.1 cm/s^2^; High SI: 403.6±46 cm/s^2^; Kolmogrov-Smirnov test; p=8.2×10^-33^; Up.: Low SI: 213.3±48.2 cm/s^2^; High SI: 598.1±76.3 cm/s^2^; p=1.9×10^-5^). (c) For each session the SI was calculated between different pairs for trials with hypermetric (overshoot; green) or hypometric (undershoot; purple) reach endpoints. Points on top correspond to time points where the two traces have significant difference (p<4×10^-5^ Bonferroni corrected for multiple testing). Inset show magnification of the difference between the two conditions in the time leading to endpoint. (d) Correlation between the sum of the SI per trial and the endpoint changes (mean±s.e.m.; − 0.25±0.02; p=2×10^-12^; t-value=10.5; dof=36; Cohen’s D= 1.7). (e) Endpoint locations for trials with the highest SI (orange), and trials with the lowest SI (blue), per cell pair (n=37 pairs; High SI: Δout: − 0.07±0.012 cm; Δup: −0.04±0.013 cm; Low SI: Δout: −0.03±0.013 cm; Δup: 0.01±0.02 cm; Wilcoxon’s signed rank; p<3.5×10^-5^; r=0.68;Wilcoxon’s signed rank). (f) The change in the outward (Out.) or vertical (Up.) endpoint position for low SI and high SI trials (Kolmogrov-Smirnov test; Δout: p-value= 1.7×10^-6^;D= 0.6; Δup: p-value= 0.11; D= 0.27; ΔEuc: Low=0.032±0.02 to high: −0.08±0.02; p-value= 1.9×10^-5^;D= 0.54).

**Extended Data Figure 5:**
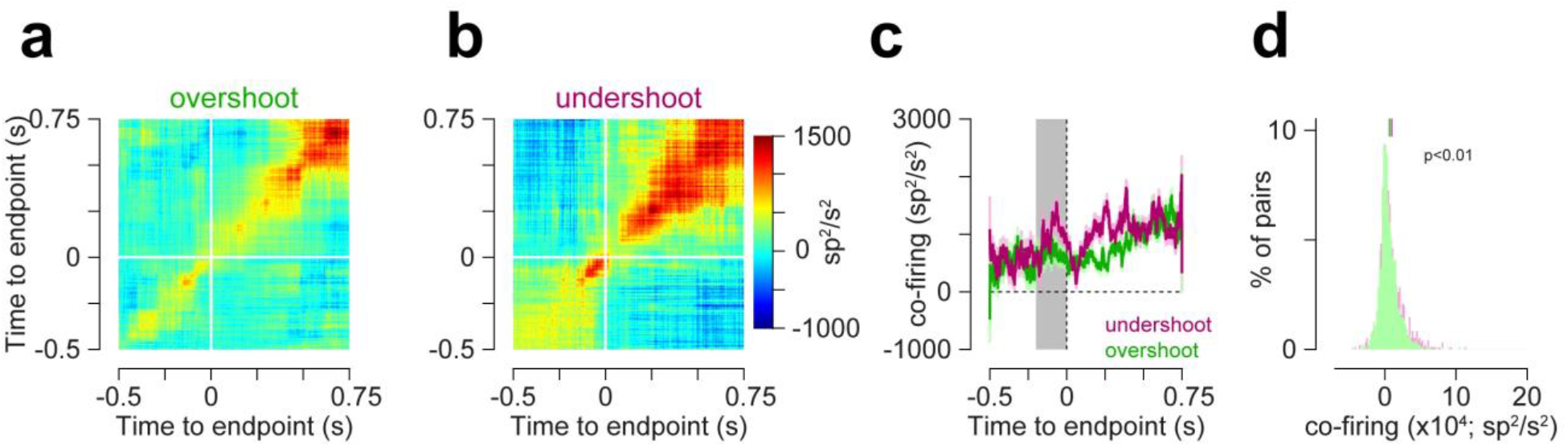
jPETHs of reaches segregated by endpoint. (a-b) jPETH matrices for the top (a) and bottom (b) quartiles of endpoint position, aligned to time of endpoint (white lines). (c) Co-firing computed along the diagonals of the jPETHs, showing average (±s.e.m.) for the hypometric (undershoot, purple) and hypermetric (overshoot, green) reaches. (d) The integral of the co-firing for the shaded area in c (Wilcoxon’s signed-rank; p=2.7×10^-7^; r=0.17).

**Extended Data Figure 6:**
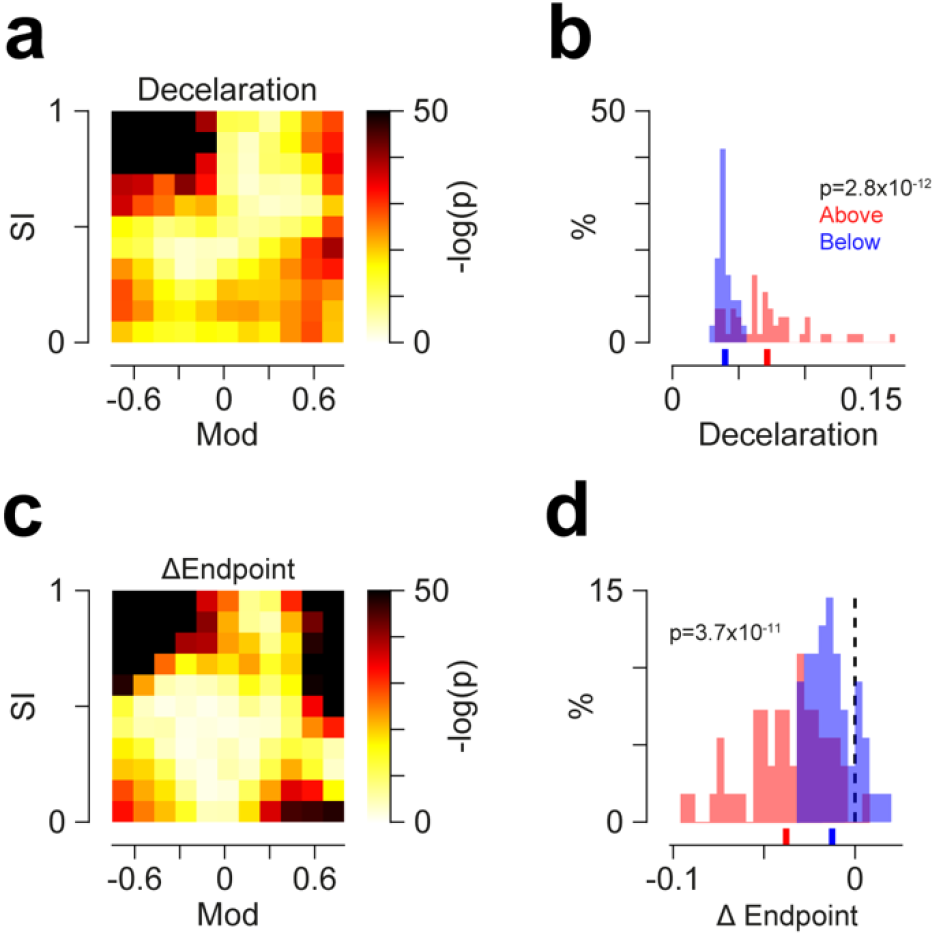
Statistics relating PC rate modulation and SI to behavioral variables (associated with data in *Fig. 3a and 3b*) (a) Surprise values (-log(p-value)) of the heatmap seen in *Fig. 3a* relative to 1,000 randomly shuffled matrices. Darker hues correspond to higher surprise and lower p-values. (b) The distribution of the bins that are located above (red) or below (blue) the main diagonal of the heatmap in 3a (under: 0.04±0.0008; above: 0.07±0.004; Wilcoxon’s ranked sum; p<7.3 x10^-12^; r= 0.92). (c) same as a), but for the data seen in *Fig. 3b*. (d) The distribution of the bins that are located above (orange) or below (blue) the main diagonal of the heatmap in 3b (under: −0.012±0.002; above: − 0.038±0.003; Wilcoxon’s ranked sum; p<9.6×10^-10^; r=0.82).

## Methods

### Animals

All procedures followed National Institutes of Health guidelines and were approved by the Institutional Animal Care and Use Committee at the University of Colorado Anschutz Medical Campus. Animals were housed in an environmentally controlled room, kept on a 12h light/dark cycle and had free access to food and water, excepting behavioral training and testing times. Adult C57BL/6 (Charles River Laboratories) mice of either sex (5 males and 2 females) were used in the experiments. The present dataset consists of all high-density recordings used in a previous study^10^. Therefore, surgical procedures and further details of task design have previously been described^10^.

### Reaching task

Briefly, after head plating, mice recovered for 2 days, then were food restricted to 80-90% of their original weight for reach training. Mice were trained to sit in a head-fixed setup, and were presented with a food pellet (20 mg, BioServ #F0163). The position of the pellet was moved further, until the mice could not retrieve it with their tongue and they began reaching for food. To encourage the animal to reach with their right hand, the pellet was positioned slightly to the right of their forelimb and a consistent position per mouse was found (∼1.2-2.5 cm from starting position). Sessions lasted for 20 successful reaches or 30 minutes. The training process lasted 15 days and mice were considered proficient once they hit 50% success rate for 3 consecutive days.

### Kinematic tracking

The position of the hand was tracked in real time using an infrared-based machine-vision motion-capture system (6 Optitrack Slim3U Cameras mounted with LED ring arrays, Motive software) at 120 frame-per-second as previously described^10,23^. The cameras were positioned in front of and to the right of the mouse and focused on the space that covered the right forelimb reach trajectory. For camera calibration and kinematic tracking, retroreflective markers (1mm) were used and affixed to the animal’s hand. To set the position and orientation of the cameras, a custom-built calibration wand and ground plane were used, in Optitrack Motive software. Motion tracking was recalibrated periodically to account for drift. The spatial origin was defined as the center of the bar the mice held during rest. Spatial blocking and camera detection threshold were adjusted to prevent erroneous tracking of other minimally infrared-reflective objects.

### Neuropixel recordings

Craniotomies were made over the cerebellum ipsilateral to the reaching arm of the mice and a custom-made recording chamber was implanted over the craniotomy. The brain was covered with triple-antibiotic ointment (GlobePharma, Inc), and the recording chamber was sealed with Qwik-sil silicone (World Precision Instruments) to facilitate multiple recording sessions. During recordings, neuropixel probes^72^ were lowered into the brain using a motorized micromanipulator (Sensapex uMp micromanipulator). Once the electrode shank spanned the putative PC layer, we waited 15 minutes for tissue to stabilize. Electrophysiology data were acquired using an OpenEphys system (https://open-ephys.org). Data were sorted offline in Kilosort2^73^ and manually curated in Phy (https://github.com/cortex-lab/phy).

### Data analysis

#### Purkinje cell identification and inclusion criteria

Units were classified as PCs or pPCs as in Calame et al. ^10^, after spike sorting. Briefly, one of two criteria were met. In the first, we identified CS, by cross correlating units with low firing rate with high-firing units and looking for the presence of CS-related simple spike pause, as well as characteristic simple spikes and CS waveforms. With our recording method, the CS was not always present in the recording. In these cases, PCs were identified based on recording location and electrophysiological criteria using the firing rate, CV2 and median absolute difference from the mediant interspike interval (MAD)^74^. Specifically, cortical units with firing rate >40 sp/s, CV2>0.2 and MAD<0.008 were labeled as PCs (see Extended Data Fig. 2 in Calame et al.^10^). This method was found to correctly identify 94% of CS-confirmed PCs^10^. Cells that were identified by either method as a PC were collectively referred to as pPCs.

To mitigate concerns that high density probes and associated sorting algorithms may inflate synchrony measurements by sampling and sorting the same unit twice, we applied a strict criterion to analyze units only if their waveforms did not overlap on the probe. To avoid double sampling of the same PC, we calculated the degree of overlap between pairs of cells and selected only PCs that did not share overlapping contacts in the 12 main contacts where they were recorded. Specifically, an overlapping index was calculated as:

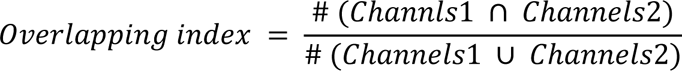

Only PCs with an overlapping index of zero were considered in the analyses reported here.

#### Joint peri-event time histogram

The joint peri-event time histogram (jPETH, Fig. 1c-e) was calculated based on previously published methods^2,25,26^. Briefly, the peri-event time histogram (PETH) was found around a specific behavioral landmark (e.g. reach endpoint). The raw jPETH matrix was calculated by inserting 1 in the (i,j)-th bin if the first neuron spiked at the i^th^ bin and the second neuron at the j^th^ bin. The jPETH predictor was calculated to correct for the rate modulation, where the (i,j)-th bin is equal to PETH1(i)*PETH(j). The jPETH was calculated in time bins of 1 ms and smoothed using two-dimensional Gaussian window of standard deviation = 3 ms.

#### Synchrony index

The synchrony index (SI) was calculated based on Sedaghat-Nejad et al^8^. Briefly, the probability of spiking in 1 ms time window was calculated for two neurons. The probability for coincident spikes in the same 1 ms time bin was computed, and the SI was defined as the ratio between the probability of coinciding spikes and the product of the probabilities of each neuron individually.

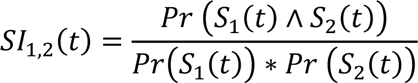

Where S1 and S2 correspond to the peri-event time histograms of unit 1 and 2.

To estimate the SI for individual trial *i* (for Fig. 2-3), we found the time-resolved co-firing for individual trial, and divided this by the time-resolved probability of each unit to fire.

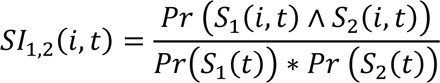

For Fig. 3, the normalized SI was calculated as follows: First, the per-trial SI was calculated. Then for each session, the trial-by-trial SI was summed over time spanning −0.3 to 0.3 s around the endpoint (spanning the entire movement) and normalized by dividing the trial-related ΣSI by the maximal ΣSI of the session. The normalization per session was made to avoid session- or mouse-related changes.

A recent critique of the SI metric showed its sensitivity to underlying firing rates when there is also positive covariance. Because we observed elevated SI preferentially in pairs which decreased their firing rates during reach, we were concerned that the metric might be misleading, artificially increasing when the denominator (i.e. firing rate) decreases. Our analyses of the diagonal of the jPETH matrix revealed that millisecond-level co-firing is indeed modulated during movement relative to shuffle controls. Thus, this structure contributes to SI along with firing rate; SI therefore reports an increase in synchronous spiking relative to the drop in firing rate which would tend to reduce their likelihood.

#### Simulating Purkinje cell populations

To test the behavior of the SI metric with simulated data, we generated spike trains for a population of Purkinje cells that decreased their firing during reach. We first constructed baseline rates of individual neurons as a Poisson process. We defined the baseline firing rate as a random integer between 50 and 110 Hz (resulting mean FR of the population∼80 Hz, closely resembling the real data baseline firing (Extending Data Fig. 3); inter spike intervals of: 9.1 to 20 ms), the firing rate during reach was set between 0 and the baseline firing rate. The dynamics and rates of the artificial firing were very similar to the real data. The onset time for “task-related” firing was set to be at 0. For each unit we simulated a session of 60 trials. We created a population of 500 units, and the SI was calculated between each unit and 5 other randomly chosen units to simulate different recording sessions. Finally, in order to synchronize the data at different levels, a specific percentage (0, 25, 50, 75 or 100%) of one unit’s spikes were randomly chosen in the time window between 0 and 0.5 s and reassigned to a shared time bins from a target unit, artificially generating a similar pattern of firing. The SI and jPETHs were then calculated based on the aforementioned methods.

### Nuclear model

To quantify the effect of PC synchrony on inhibitory current integration dynamics in postsynaptic nuclear cells, we built a model of nuclear cells receiving a converging input of 40 PCs (Fig. 4). The firing rate of the PCs ranged between 20 sp/s to 120 sp/s, in 2 sp/s steps. The firing of PCs was constructed for 1,000 time points (corresponding to 1 s) from a random Poisson distribution of the interspike intervals (8.33 ms to 50 ms). For each frequency, we used variable synchrony level (0, 1, 2,….,60%) and the model was run for 1,000 iterations. To quantify how the synchrony levels change the level of inhibition on the nuclear neuron we calculated the following:

a. Arriving PC inputs (Fig. 4c,f and h): This corresponds to the product of the firing rate in 1 bin across the 40 PCs, such that if 1 PC fires at a time i, the nuclear cells will receive a PC spike at that time. Importantly, if more than one PC fires at the same time bin (synchronized), the response of the nuclear cell will be as though just one PC fired, owing to the overlap of the fast (τ = 2.5ms) inhibition of each PC on the nuclear cell.
b. Inter-spike interval (ISI, Fig. 4i): This corresponds to the ISI of the arriving PC spikes.
c. Inhibitory current (Fig. 4d,g,j and k): PC spike arrivals (from one PC or more) induced IPSCs in the postsynaptic nuclear cell modeled as a skewed gaussian with a decay parameter of 2.5 ms, as measured at physiological temperatures (Person and Raman 2012a), and IPSCs temporally summated within that decay window.

### Acute brain slice preparation

#### Tissue Preparation

Postnatal day 32-42 C57BL/6 mice of both sexes (n=15 mice, 7 females 8 males) were deeply anesthetized with isoflurane before transcardial perfusion with warmed Tyrode’s solution containing (in mM) 123.75 NaCl, 26 NaHCO_3_, 10 glucose, 3.5 KCl, 1.25 NaH_2_PO_4_, 1.5 CaCl_2_, and 1 MgCl_2_. Solutions were oxygenated with 95% O_2_, 5% CO_2_ carbogen. Following perfusion, tissue was harvested and 300 µm thick parasagittal slices were made using a Leica VT1000 S vibratome (Leica, Wetzlar, Germany). Slices were incubated at 34°C for 1 hour before being held at room temperature for the duration of the experiments (approx. 5 hours).

#### Data Acquisition

Slices were transferred to a recording chamber and were continuously perfused with oxygenated Tyrode’s solution at 32°C - 34°C which included synaptic blockers SR95531 (gabazine, 10µM), 6,7-dinitroquinoxaline-2,3-dione (DNQX, 10µM), and (R)-CPP (10µM). All drugs were purchased from Tocris (Bristol, United Kingdom). Glass patch pipettes (Sutter Instruments BF150-86-7.5) were pulled using a Sutter P97 (Sutter Instrument, Novato, CA, USA) to produce tip resistances of 2 – 4 MΩ and were filled with a K-gluconate intracellular solution (290 mOsm, pH 7.37) consisting of 132 K-gluconate, 4.4 Na-gluconate, 4.4 NaCl, 2.2 MgCl_2_, 1.1 EGTA, 11 HEPES, 22 sucrose, 14 TRIS creatine phosphate, 4 Mg-ATP, 0.3 and TRIS-GTP. Stable whole-cell recordings were acquired at 100 kHz in pClamp 10.3 and pClamp 10.7 (Molecular Devices, San Jose, CA, USA) via an Axon 700B multiclamp amplifier and Digidata 1440 digitizer. Cells were visually identified and included for recording based on an access resistance below 20MΩ, and the demonstration of a robust fast inward sodium current in voltage clamp and overshooting action potentials in current clamp.

#### Dynamic Clamp

Dynamic clamp experiments were conducted based on hardware and software designs described previously^75^. IPSGs were simulated with characteristics described by(Person and Raman 2012a), specifically, 5 nS peak conductance with a 100 µsec rise time and 2.5 msec decay time. The 100 kHz sampling rate used allowed us to simulate synaptic integration at instantaneous rates of up to 50 kHz and thus allowed us to model a convergent Purkinje cell pool of up to 50 cells. Purkinje cell trains were constructed based on previously published PC population firing rates during reaching movements^10^. Briefly, interspike intervals of a PC “seed” trace were established to reproduce a PETH of from a given reach velocity range (0-20%; 60-80% and 80-100% peak speed). Spike times for other 39 additional simulated PCs within that “pool” were then drawn from seed trace spike times with the addition of random jitter drawn from a normal distribution. This method allowed us to modulate population synchrony dynamically throughout the reach epoch by modulating the standard deviation of these distributions, while maintaining the net time-varying population activity observed in Calame et al^10^. Finally, for experiments modulating PC rate and synchrony, PC pools with mean rates of 40-100 Hz in 10 Hz intervals were generated. Within these pools synchrony was swept from an SI of 1 to 2 throughout a 4 second epoch.

#### Statistical analysis

The shuffled data (in Fig. 1g-i, Extended Data Fig. 1d-f, Extended Data Figure 2) were created by permuting/randomizing the order of the recorded trials, such that the average firing rate of each cell was kept, however, the trial-to-trial variability between the cells is impaired. This method of shuffling was chosen (as opposed to spike times shuffling, for example) given recent critique of the SI method, where it was suggested that the SI is highly affected by the firing rate modulation. To test for significant differences in time series data, (in Extended data Figure 2 and 4), for each time bin (1 ms and non overlapping), the values that were found in one group (in Extended Data Fig. 2a for example, 951 values for real) were tested against the values in the other group (951 values for shuffled data), using running paired Student’s t tests. Only p values smaller than 0.00004 were considered significant, after correcting for multiple testing (Bonferroni correction for the number of time bins: 0.05/1250=0.00004).This method of testing the difference between two time series, in time, is similar to methods used in the past^76–79^.

Data presented in the manuscript treat each neuron as an independent sample. All data were tested for normality using Kolmogrov-Smirnov test to choose the proper statistical test. All t-tests or Wilcoxon’s signed rank are paired, unless stated otherwise. In box plots, the box displays the median and the 25th and 75th percentiles, the lines extend to the 10th and 90th percentiles of the data.

Effect sizes were estimated using Cohen’s d, for parametric tests. For non-parametric tests, the effect size (r) was calculated as the z-statistics divided by the square root of the sample size. No statistical tests were used to predetermine required sample size. Randomizations and controls are described in the main text and methods. The experimenters were not blind to conditions during experiments and outcome assessments. Nonparametric tests were used when the dataset violated normality assumptions. Multiple regression analysis was performed to test how both synchrony and firing rate modulation together affect movement kinematics. Multiple linear regression was computed on the matrices seen in Fig. 3 using the Matlab function *fitlm*. The input to the function consisted of the kinematics data in a bin-by-bin fashion (response variable), and the SI or modulations corresponding to each bin (predictors).

All data analyses were made in Matlab (Mathworks, 2020). Raw data and some waveform analysis were made in Python and Visual Studio. Original scripts are available at: https://github.com/AbedNashef/Sychrony_paper

## Acknowledgements

We would like to thank Dr. Daniel Denman for his help with waveform analysis, and Drs. David DiGregorio and Jason Christie and members of the Person lab for their insightful comments on the manuscript. We thank Maham Haq for participating in the initial phases of the study. This study was supported by the Life Sciences Research Foundation (and Amgen) and the Edmond and Lily Safra Campus Post-doctoral fellowship to AN; F31-NS113395 to DJC; and NS114430 and NSF CAREER to ALP.

## Author Contributions

AN and ALP conceptualized the study. DJC acquired the neural data. MSS performed the *in vitro* experiments and analyzed the in vitro data. AN analyzed the in vivo data and constructed models. AN and ALP wrote the paper.

## List of References

1. Bastian, A. J., Martin, T. A., Keating, J. G. & Thach, W. T. Cerebellar ataxia: Abnormal control of interaction torques across multiple joints. J. Neurophysiol. 76, 492–509 (1996).

2. Nashef, A., Mitelman, R., Harel, R., Joshua, M. & Prut, Y. Area-specific thalamocortical synchronization underlies the transition from motor planning to execution. Proc. Natl. Acad. Sci. U. S. A. 118, (2021).

3. Nashef, A., Cohen, O., Harel, R., Israel, Z. & Prut, Y. Reversible Block of Cerebellar Outflow Reveals Cortical Circuitry for Motor Coordination. Cell Rep. 27, 2608–2619.e4 (2019).

4. Nashef, A., Cohen, O., Israel, Z., Harel, R. & Prut, Y. Cerebellar Shaping of Motor Cortical Firing Is Correlated with Timing of Motor Actions. Cell Rep. 23, 1275–1285 (2018).

5. Gao, Z. et al. A cortico-cerebellar loop for motor planning. Nature 563, 113–116 (2018).

6. Lee, J. H. et al. Cerebellar granule cell signaling is indispensable for normal motor performance. Cell Rep. 42, 112429 (2023).

7. Herzfeld, D. J., Joshua, M. & Lisberger, S. G. Rate versus synchrony codes for cerebellar control of motor behavior. bioRxiv (2023) doi:10.1101/2023.02.17.529019.

8. Sedaghat-Nejad, E., Pi, J. S., Hage, P., Fakharian, M. A. & Shadmehr, R. Synchronous spiking of cerebellar Purkinje cells during control of movements. Proc. Natl. Acad. Sci. U. S. A. 119, (2022).

9. Bell, C. C. & Grimm, R. J. Discharge properties of Purkinje cells recorded on single and double microelectrodes. J. Neurophysiol. 32, 1044–55 (1969).

10. Calame, D. J., Becker, M. I. & Person, A. L. Cerebellar associative learning underlies skilled reach adaptation. Nat. Neurosci. 26, 1068–1079 (2023).

11. Coltz, J. D., Johnson, M. T. V. & Ebner, T. J. Cerebellar Purkinje Cell Simple Spike Discharge Encodes Movement Velocity in Primates during Visuomotor Arm Tracking. J. Neurosci. 19, 1782 (1999).

12. Heck, D. H., Thach, W. T. & Keating, J. G. On-beam synchrony in the cerebellum as the mechanism for the timing and coordination of movement. Proc. Natl. Acad. Sci. U. S. A. 104, 7658–7663 (2007).

13. Herzfeld, D. J., Kojima, Y., Soetedjo, R. & Shadmehr, R. Encoding of action by the Purkinje cells of the cerebellum. Nature 526, 439–441 (2015).

14. Hewitt, A. L., Popa, L. S., Pasalar, S., Hendrix, C. M. & Ebner, T. J. Representation of limb kinematics in Purkinje cell simple spike discharge is conserved across multiple tasks. J. Neurophysiol. 106, 2232 (2011).

15. Person, A. L. & Raman, I. M. Purkinje neuron synchrony elicits time-locked spiking in the cerebellar nuclei. Nature 481, 502–505 (2012).

16. Roitman, A. V., Pasalar, S., Johnson, M. T. V. & Ebner, T. J. Position, Direction of Movement, and Speed Tuning of Cerebellar Purkinje Cells during Circular Manual Tracking in Monkey. J. Neurosci. 25, 9244 (2005).

17. de Solages, C. et al. High-Frequency Organization and Synchrony of Activity in the Purkinje Cell Layer of the Cerebellum. Neuron 58, 775–788 (2008).

18. Wise, A. K., Cerminara, N. L., Marple-Horvat, D. E. & Apps, R. Mechanisms of synchronous activity in cerebellar Purkinje cells. J. Physiol. 588, 2373–2390 (2010).

19. Ohtsuka, K. & Noda, H. Discharge properties of Purkinje cells in the oculomotor vermis during visually guided saccades in the macaque monkey. https://doi.org/10.1152/jn.1995.74.5.1828 74, 1828–1840 (1995).

20. Noda, H., Murakami, S., Yamada, J., Tamaki, Y. & Aso, T. Saccadic eye movements evoked by microstimulation of the fastigial nucleus of macaque monkeys. https://doi.org/10.1152/jn.1988.60.3.1036 60, 1036–1052 (1988).

21. Spencer, R. M. C., Zelaznik, H. N., Diedrichsen, J. & Ivry, R. B. Disrupted timing of discontinuous but not continuous movements by cerebellar lesions. Science (80-. ). 300, 1437–1439 (2003).

22. Bo, J., Block, H. J., Clark, J. E. & Bastian, A. J. A cerebellar deficit in sensorimotor prediction explains movement timing variability. J. Neurophysiol. 100, 2825–2832 (2008).

23. Becker, M. I. & Person, A. L. Cerebellar Control of Reach Kinematics for Endpoint Precision. Neuron 103, 335–348.e5 (2019).

24. Lee, K. H. et al. Circuit Mechanisms Underlying Motor Memory Formation in the Cerebellum. Neuron 86, 529–540 (2015).

25. Gerstein, G. L. & Perkel, D. H. Simultaneously Recorded Trains of Action Potentials: Analysis and Functional Interpretation. Science (80-. ). 164, 828–830 (1969).

26. Aertsen, A. M., Gerstein, G. L., Habib, M. K. & Palm, G. Dynamics of neuronal firing correlation: modulation of ‘effective connectivity’. J. Neurophysiol. 61, 900–917 (1989).

27. Ebner, T. J. & Bloedel, J. R. Correlation Between Activity of Purkinje Cells and Its Modification by Natural Peripheral Stimuli. J. Neurophysiol. 45, (1981).

28. Medina, J. F. & Lisberger, S. G. Variation, Signal, and Noise in Cerebellar Sensory– Motor Processing for Smooth-Pursuit Eye Movements. J. Neurosci. 27, 6832–6842 (2007).

29. Sauerbrei, B. A., Lubenov, E. V. & Siapas, A. G. Structured Variability in Purkinje Cell Activity during Locomotion. Neuron 87, 840–852 (2015).

30. Baker, S. N., Kilner, J. M., Pinches, E. M. & Lemon, R. N. The role of synchrony and oscillations in the motor output. Exp. Brain Res. 128, 109–117 (1999).

31. Fetz, E. E. & Cheney, P. D. Postspike facilitation of forelimb muscle activity by primate corticomotoneuronal cells. J. Neurophysiol. 44, 751–772 (1980).

32. Person, A. L. & Raman, I. M. Synchrony and neural coding in cerebellar circuits. Front. Neural Circuits (2012) doi:https://doi.org/10.3389/fncir.2012.00097.

33. Herzfeld, D. J., Kojima, Y., Soetedjo, R. & Shadmehr, R. Encoding of error and learning to correct that error by the Purkinje cells of the cerebellum. Nat. Neurosci. 21, 736–743 (2018).

34. Yang, Y. & Lisberger, S. G. Purkinje-cell plasticity and cerebellar motor learning are graded by complex-spike duration. Nat. 2014 5107506 510, 529–532 (2014).

35. Shin, S. L. & De Schutter, E. Dynamic synchronization of purkinje cell simple spikes. J. Neurophysiol. 96, 3485–3491 (2006).

36. Bell, C. C. & Kawasaki, T. Relations Among Climbing Fiber Responses of Nearby Purkinje Cells. J. Neurophysiol. 35, 155–169 (1971).

37. Bosman, L. W. J. et al. Encoding of whisker input by cerebellar Purkinje cells. J. Physiol. 588, 3757–3783 (2010).

38. Brown, S. T. & Raman, I. M. Sensorimotor Integration and Amplification of Reflexive Whisking by Well-Timed Spiking in the Cerebellar Corticonuclear Circuit. Neuron 99, 564–575.e2 (2018).

39. Han, K. S. et al. Ephaptic Coupling Promotes Synchronous Firing of Cerebellar Purkinje Cells. Neuron 100, 564–578.e3 (2018).

40. Turecek, J. & Regehr, W. G. Cerebellar and vestibular nuclear synapses in the inferior olive have distinct release kinetics and neurotransmitters. Elife 9, 1–22 (2020).

41. De Zeeuw, C. I. Bidirectional learning in upbound and downbound microzones of the cerebellum. Nat. Rev. Neurosci. 22, 92–110 (2020).

42. Gao, W., Chen, G., Reinert, K. C. & Ebner, T. J. Cerebellar Cortical Molecular Layer Inhibition Is Organized in Parasagittal Zones. J. Neurosci. 26, 8377–8387 (2006).

43. Diño, M. R., Willard, F. H. & Mugnaini, E. Distribution of unipolar brush cells and other calretinin immunoreactive components in the mammalian cerebellar cortex. J. Neurocytol. 28, 99–123 (1999).

44. Ekerot, C. F., Jörntell, H. & Garwicz, M. Functional relation between corticonuclear input and movements evoked on microstimulation in cerebellar nucleus interpositus anterior in the cat. Exp. Brain Res. 106, 365–376 (1995).

45. Low, A. Y. T. et al. Precision of Discrete and Rhythmic Forelimb Movements Requires a Distinct Neuronal Subpopulation in the Interposed Anterior Nucleus. Cell Rep. 22, 2322– 2333 (2018).

46. Jackson, A., Gee, V. J., Baker, S. N. & Lemon, R. N. Synchrony between neurons with similar muscle fields in monkey motor cortex. Neuron 38, 115–125 (2003).

47. Vaadia, E. et al. Dynamics of neuronal interactions in monkey cortex in relation to behavioral events. Nature vol. 373 515–518 (1995).

48. Dan, Y., Alonso, J. M., Usrey, W. M. & Reid, R. C. Coding of visual information by precisely correlated spikes in the lateral geniculate nucleus. Nat. Neurosci. 1, 501–507 (1998).

49. Hatsopoulos, N. G., Ojakangas, C. L., Paninski, L. & Donoghue, J. P. Information about movement direction obtained from synchronous activity of motor cortical neurons. Proc. Natl. Acad. Sci. 95, 15706–15711 (1998).

50. Stark, E., Globerson, A., Asher, I. & Abeles, M. Correlations between Groups of Premotor Neurons Carry Information about Prehension. J. Neurosci. 28, 10618–10630 (2008).

51. Bruno, R. M. & Sakmann, B. Cortex Is Driven by Weak but Synchronously Active Thalamocortical Synapses. Science (80-. ). 312, 1622–1627 (2006).

52. Gauck, V. & Jaeger, D. The Control of Rate and Timing of Spikes in the Deep Cerebellar Nuclei by Inhibition. J. Neurosci. 20, 3006–3016 (2000).

53. Wu, Y., Raman, I. M., Raman, I. M. & Wu, Y. Facilitation of mossy fibre-driven spiking in the cerebellar nuclei by the synchrony of inhibition. J. Physiol. 595, 5245–5264 (2017).

54. Wu, S., Wardak, A., Khan, M. M., Chen, C. H. & Regehr, W. G. Implications of variable synaptic weights for rate and temporal coding of cerebellar outputs. bioRxiv 2023.05.25.542308 (2023) doi:10.1101/2023.05.25.542308.

55. Sarnaik, R. & Raman, I. M. Control of voluntary and optogenetically perturbed locomotion by spike rate and timing of neurons of the mouse cerebellar nuclei. Elife 7, (2018).

56. Kim, J. & Augustine, G. J. Molecular Layer Interneurons: Key Elements of Cerebellar Network Computation and Behavior. Neuroscience 462, 22–35 (2021).

57. Blot, A. & Barbour, B. Ultra-rapid axon-axon ephaptic inhibition of cerebellar Purkinje cells by the pinceau. Nat. Neurosci. 17, 289–295 (2014).

58. Hoehne, A., McFadden, M. H. & Digregorio, D. A. Feed-forward recruitment of electrical synapses enhances synchronous spiking in the mouse cerebellar cortex. Elife 9, 1–26 (2020).

59. Arlt, C. & Häusser, M. Microcircuit Rules Governing Impact of Single Interneurons on Purkinje Cell Output In Vivo. Cell Rep. 30, 3020–3035.e3 (2020).

60. Blot, A. et al. Time-invariant feed-forward inhibition of Purkinje cells in the cerebellar cortex in vivo. J. Physiol. 594, 2729–2749 (2016).

61. Nashef, A., Cohen, O., Perlmutter, S. I. & Prut, Y. A cerebellar origin of feedforward inhibition to the motor cortex in non-human primates. Cell Rep. 39, (2022).

62. Pouille, F. & Scanziani, M. Enforcement of Temporal Fidelity in Pyramidal Cells by Somatic Feed-Forward Inhibition. Science (80-. ). 293, 1159–1163 (2001).

63. Gaffield, M. A., Rowan, M. J. M., Amat, S. B., Hirai, H. & Christie, J. M. Inhibition gates supralinear Ca2+ signaling in purkinje cell dendrites during practiced movements. Elife 7, (2018).

64. Urrutia-Piñones, J., Morales-Moraga, C., Sanguinetti-González, N., Escobar, A. P. & Chiu, C. Q. Long-Range GABAergic Projections of Cortical Origin in Brain Function. Front. Syst. Neurosci. 16, 841869 (2022).

65. Wichmann, T. & Dostrovsky, J. O. Pathological basal ganglia activity in movement disorders. Neuroscience 198, 232–244 (2011).

66. Villalobos, C. A. & Basso, M. A. Optogenetic activation of the inhibitory nigro-collicular circuit evokes contralateral orienting movements in mice. Cell Rep. 39, 110699 (2022).

67. Howe, M. W., Atallah, H. E., McCool, A., Gibson, D. J. & Graybiel, A. M. Habit learning is associated with major shifts in frequencies of oscillatory activity and synchronized spike firing in striatum. Proc. Natl. Acad. Sci. U. S. A. 108, 16801–16806 (2011).

68. White, J. J. & Sillitoe, R. V. Genetic silencing of olivocerebellar synapses causes dystonia-like behaviour in mice. Nat. Commun. 8, 1–16 (2017).

69. Sausbier, M. et al. Cerebellar ataxia and Purkinje cell dysfunction caused by Ca2+-activated K+channel deficiency. Proc. Natl. Acad. Sci. U. S. A. 101, 9474–9478 (2004).

70. Walter, J. T., Alviña, K., Womack, M. D., Chevez, C. & Khodakhah, K. Decreases in the precision of Purkinje cell pacemaking cause cerebellar dysfunction and ataxia. Nat. Neurosci. 9, 389–397 (2006).

71. Brown, A. M. et al. Purkinje cell misfiring generates high-amplitude action tremors that are corrected by cerebellar deep brain stimulation. Elife 9, (2020).

72. Jun, J. J. et al. Fully integrated silicon probes for high-density recording of neural activity. Nature 551, 232–236 (2017).

73. Pachitariu, M., Steinmetz, N., Kadir, S., Carandini, M. & D., H. K. Kilosort: realtime spike-sorting for extracellular electrophysiology with hundreds of channels. bioRxiv 061481 (2016) doi:10.1101/061481.

74. Hensbroek, R. A. et al. Identifying Purkinje cells using only their spontaneous simple spike activity. J. Neurosci. Methods 232, 173–180 (2014).

75. Desai, N. S., Gray, R. & Johnston, D. A Dynamic Clamp on Every Rig. eNeuro 4, (2017).

76. Kong, E., Lee, K.-H., Do, J., Kim, P. & Lee, D. Dynamic and stable hippocampal representations of social identity and reward expectation support associative social memory in male mice. Nat. Commun. 14, 1–20 (2023).

77. Markanday, A., Hong, S., Inoue, J., De Schutter, E. & Thier, P. Multidimensional cerebellar computations for flexible kinematic control of movements. Nat. Commun. 14, 1–16 (2023).

78. Catz, N., Dicke, P. W. & Thier, P. Cerebellar-dependent motor learning is based on pruning a Purkinje cell population response. Proc. Natl. Acad. Sci. U. S. A. 105, 7309–7314 (2008).

79. Ilg, U. J., Schumann, S. & Thier, P. Posterior Parietal Cortex Neurons Encode Target Motion in World-Centered Coordinates. Neuron 43, 145–151 (2004).

